# A Universal, AI-based Design Framework for Efficient Manufacturing of mRNA Therapeutics

**DOI:** 10.64898/2026.03.09.710039

**Authors:** Kuo-Chieh Liao, Giuseppe Maccari, Giorgio Ciano, Roland Huber, Tobias von der Haar, Cheng-Yong Tham, Nicholas Ting Xun Ong, Paola Florez de Sessions, Tzuen Yih Saw, Ting Wei Lim, Craig Martin, Mark Dickman, Zoltan Kis, Harris Makatsoris, Alexander van Asbeck, Yue Wan, Duccio Medini

## Abstract

The growth of mRNA therapeutics is limited by bespoke manufacturing processes. To overcome this barrier to access and innovation, we introduce an AI-driven framework that decouples sequence design from manufacturing, analogous to the universal design principles that revolutionized the semiconductor industry. We performed a large-scale screen, quantifying the in vitro transcription (IVT) efficiency of one million diverse sequences. We then trained an interpretable deep learning model that accurately predicts manufacturability from sequences alone and learned underlying molecular mechanisms. An algorithm using this model prospectively improved the IVT yield of a vaccine and a gene-editing therapeutic by over 7.5-fold. Co-optimization for manufacturability and translation efficiency provided improvements over state-of-the-art commercial mRNA products. Our AI-driven framework establishes a universal design paradigm with the promise to democratize and accelerate the development of mRNA medicines, potentially unlocking a new era in biotechnology.

## Main text

COVID-19 vaccines demonstrated the transformative potential of mRNA therapeutics,^1^ which enable an organism to produce a desired protein by delivering genetic information to host cells. This concept can be applied to many health conditions^2–8^. Diverse mRNA designs can encode for antigens^9^, monoclonal antibodies^10^, hormones^11^, gene therapy complexes^12^, and in-vivo chimeric antigen receptor T cell therapies^13^. However, product-specific manufacturing requirements hamper the potential development of a multi-product mRNA platform.

Like all biologic products, each mRNA therapeutic requires a specific manufacturing process, and scale-ups costing hundreds of millions of dollars over several years^14–15^. The primary RNA manufacturing step is *in vitro* transcription (IVT)^16^, which uses an RNA polymerase enzyme to transcribe DNA into the desired mRNA molecule^17^. Establishing and optimizing IVT is also time-intensive and expensive, with design-dependent yield and quality^18–19^. The semiconductor industry solved a similar problem by developing universal design rules for very large-scale integration, decoupling product design from manufacturing process specifications and creating an industry with sustained growth^14–15, 20^. Following this idea, we sought to create a design framework to ensure the manufacturability of mRNA products at high yield and quality, independent of the specific IVT process adopted.

The key to ensuring robust, timely mRNA production is DNA templates that can make large amounts of high-quality mRNA without tailored process adaptations. However, current mRNA product design efforts focus on codon optimization to improve mRNA translatability in the cell^21–22^. The presence of canonical RNA polymerase terminators^23^, abundant secondary structure elements, and poly-purine/pyrimidine rich sequences all influence IVT^24–25^. Attempts to improve manufacturing efficiency by engineering the T7 polymerase enzyme have been primarily geared towards reducing byproducts such as dsRNA or loop-back RNAs^26–27^.

During the COVID-19 pandemic, IVT processes were developed to large scale using batch process design^28–30^, but are now evolving towards more scalable, standardizable approaches, including semi-continuous and continuous flow^31–33^. Therefore, we sought to identify requirements to ensure the design of DNA templates that achieve high yields and high quality IVT-derived mRNAs using diverse processes. (Fig. 1).

**Figure 1:**
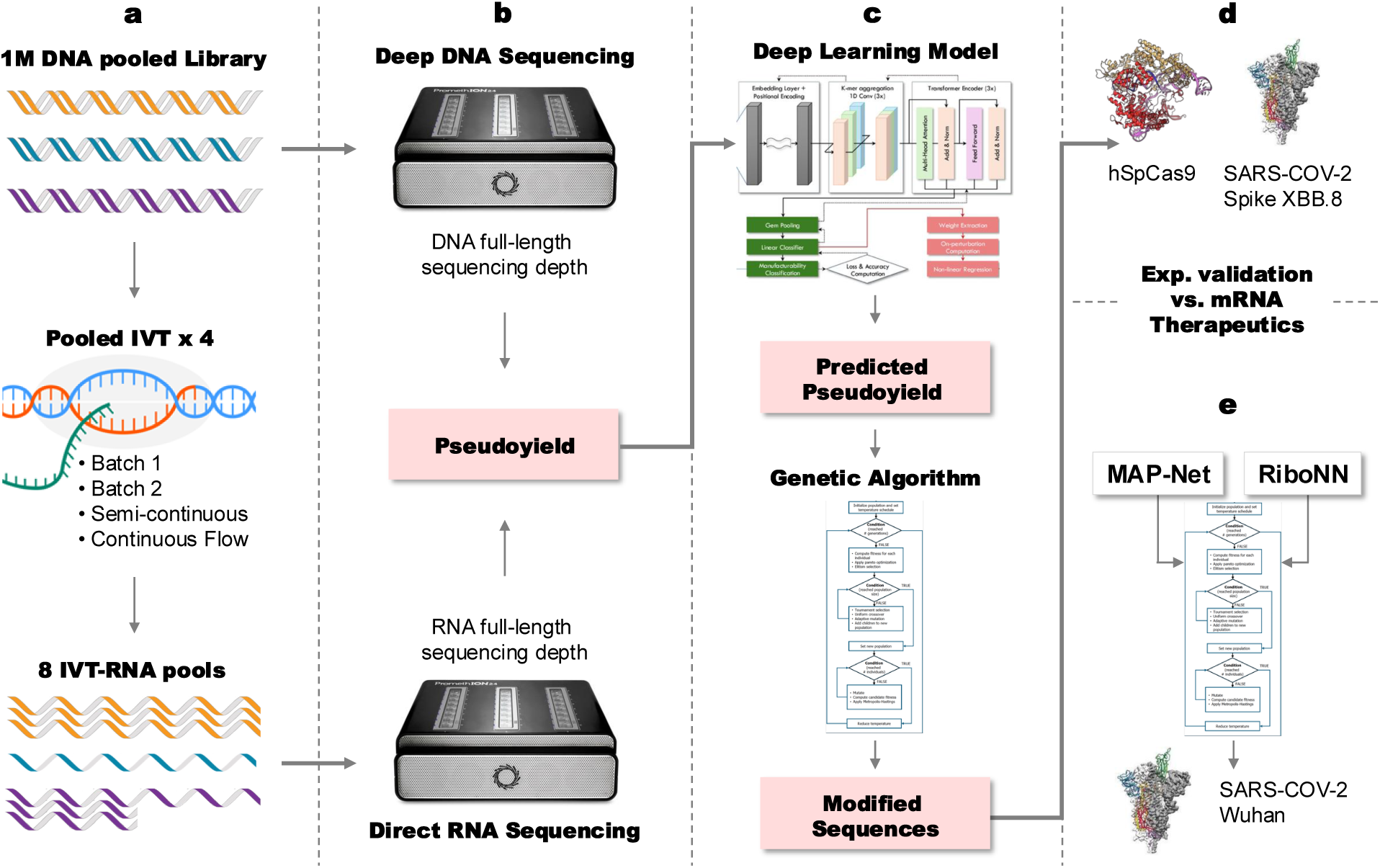
Key steps in developing a universal framework for manufacturable mRNA therapeutics. **A.** A DNA library of 1 million templates that contain 100% of all possible 11-mers is used as input for pooled in vitro transcription (IVT) using different manufacturing processes to create eight IVT-RNA pools. **B.** Oxford Nanopore Technology Sequencing is used to quantify full-length, (untruncated) DNA and IVT-RNA templates (3 and 8 pools, respectively). The sequencing depths generated are used to calculate a pseudoyield (PY), representative of the amount of full-length (untruncated) IVT-RNA produced per DNA template. **C.** PY values from the 1M DNA library are used to train, validate and test a Multi-scale Attention Projection Network (MAP-Net), that accurately predicts the PY of unknown DNA templates. MAP-Net is then used within a Genetic Algorithm that generates new synonymously mutated templates with different predicted PY for any DNA template. **D.** The MAP-Net-based genetic algorithm is used on DNA templates for two real-world mRNA products, hSpCas9 and XBB.8 Spike, generating for each 4 synonymous templates with predicted PYs higher (2) and lower (2) than the starting template. Yields measured for full-length transcripts in individual IVT experiments validate MAP-Net PY predictions. **E**. The same Genetic Algorithm is now used with MAP-Net and RiboNN as fitness functions to co-optimize the Wuhan Spike sequence for translatability and manufacturability, and results are compared with commercial SARS-COV-2 vaccine sequences.

## Results

### Long-read, direct DNA and RNA sequencing quantifies a million templates pre- and post-IVT

We designed a library of 1 million 300-mers comprising genomic sequences from five kingdoms, including all coding sequences in the human transcriptome, and sequences from major bacterial and viral species (Fig. 1A, Fig. S1A). Each 300-mer included 26 bases of the T7 promoter, 254 bases of 1M diverse sequences, and 20 adenines to enable downstream IVT and deep sequencing (Fig. S1B). The sequences covered 100% of all possible 11-mers and 95% of 12-mers and were filtered for <90% sequence similarity for maximal diversity (Fig. S1C, Methods 1).

We used PCR amplification to produce the final double-stranded DNA template library for downstream IVT (Methods 2). This was then analyzed in three technical replicates using DNA nanopore sequencing (Methods 3.1). We detected 97.8% of the initially designed templates and obtained a high correlation of sequencing depth per template (*r* > 0.97, Fig. S2A). After filtering for full-length sequencing reads (Fig. S2A) and for a minimum sequencing depth of 0.2 counts per million (CPM, Fig. S2C), the total library was 780,832 full-length DNA templates with sufficient abundance for downstream analysis (Fig. S2D). The median coefficient of variation of the CPM was 8.3%, and after filtering, no templates showed variations >30%, indicating good reproducibility (Fig. S2E).

Since RNA yields may vary with different manufacturing processes, we used four IVT processes in two biological replicates from the same dsDNA library to produce *in vitro* transcripts from the PCR-amplified template pool (Fig. 1). The processes were two batch conditions—one commercial kit and one process using in-house materials—semi-continuous and continuous flow (Methods 2.1). We purified the resulting eight IVT-RNA pools and performed direct RNA sequencing to determine the relative abundance of mRNAs in each library (Methods 3.1, Table S2).

Sequencing generated 50.6 to 64.0 million reads per IVT-process/replicate, with a high correlation within (*r* >0.97) and between IVT processes (*r* >0.95, Fig. S3A), suggesting processes robustness and comparability. Selecting full-length reads (Fig. S3B) we obtained 47.2 to 59.6 million full-length IVT-RNAs per IVT-process/replicate, or 93-94% of the whole reads per sample. The median variation within-process of the sequencing depth per template ranged from 7.5% to 8.5%, indicating good reproducibility of the direct RNA sequencing method as a means to quantify full-length IVT-RNAs in a pool (Fig. S3C).

### mRNA manufacturability varies over 100-fold between different templates

To determine the number of full-length IVT-RNAs transcribed per DNA template molecule, we calculated a pseudoyield (PY) value by normalizing, for each template, the sequencing depth of full-length IVT-RNAs to that of full-length DNA (Methods 4.2). Since native DNA and RNA molecules were directly sequenced without amplification, PY represents a quantitative measurement of RNAs transcribed from each specific DNA template without PCR bias. All IVT processes produced similar PY distributions (Fig. 2), and we calculated a consensus PY across all manufacturing runs. We observed a unimodal distribution whereby 67% of the templates show consensus PYs between 0.834 and 1.225, or 1.03 ± 19% (Fig. 2A); this range is compatible with the experimental variability measured for the PY values (median CV ≃20%, Fig. 2A). Within this central PY range, templates followed a linear trend whereby DNA and IVT-RNA amounts were directly proportional, within <1.5 fold of each other (Fig. 2B). Outside this range, two sixths of the library (16.5% each, 260,277 templates total) showed high/low PYs, with DNA templates producing as high as 16 times more full-length mRNA than expected based on DNA abundance, or as low as zero full-length IVT-RNA output, indicating high and low manufacturability, respectively.

**Figure 2:**
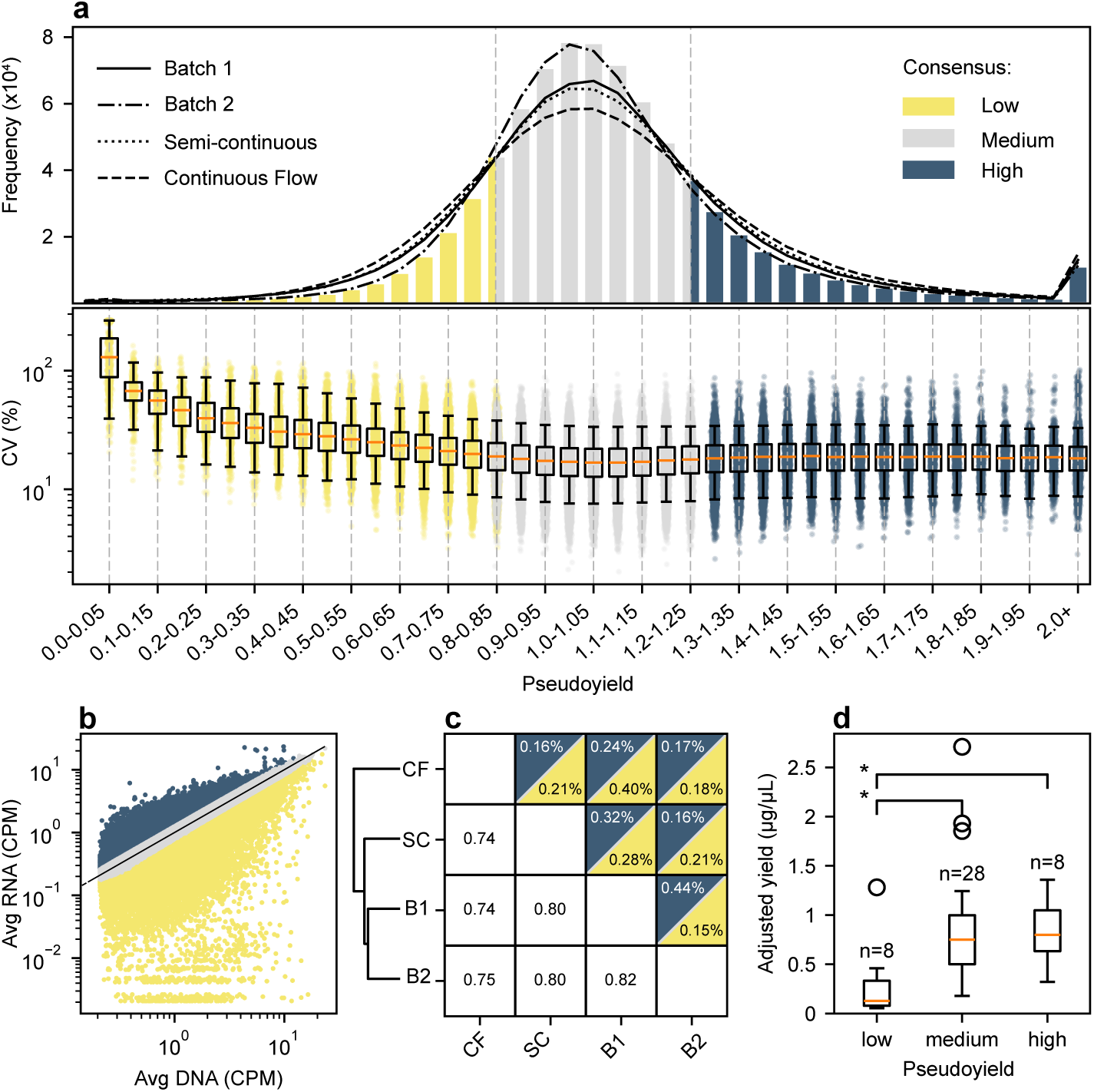
Pseudoyields for the 1Million DNA template library from 4 IVT processes. **A.** Top panel: distribution of PYs calculated according to Supplementary Methods 4.2, for each IVT process (lines) and for the consensus PY (bar plot). Dashed vertical lines limit the lower/higher sextile of the PY distribution: Low <0.835 (yellow), High >1.25 (blue), Medium otherwise (gray). Bottom panel: Coefficients of variation (CV) for consensus PYs of each template in the library are shown grouped in PY bins of 0.05 width: individual templates (dots) and bin distributions (boxplots). **B.** Scatterplot of sequencing counts per million (CPM) values from DNA templates (x-axis, average of 3 DNA sequencing runs) and IVT-RNAs (y-axis, average of 8 IVT runs). Color coding as in Panel A. Solid curve is an y=x guide to the eye. **C.** Lower-left triangle: Pearson correlation coefficient between each pair of IVT processes, considering all DNA templates. Upper-right triangle: percentage of templates classified as high-yield from the left/bottom process and low-yield from the bottom/left process (yellow/blue, respectively). The dendrogram represents the agglomerative clustering results generated from the PY correlation **D.** Boxplot of measured yields, adjusted to account only for full-length products (Supplementary Methods 2.2), from 44 randomly selected templates, grouped by low-, medium- and high-predicted PY. Medians are shown in orange, black boxes represent interquartile ranges (IQRs), whiskers are 1.5*IQR, outliers as circles. Horizontal brackets denote significant differences between low- *vs.* high- and medium- *vs.* low- (Bonferroni-corrected Mann Whitney U test p < 0.01).

PYs from the batch process conditions were closer to those from the semi-continuous than to those from continuous flow IVT (Fig. 2C). Overall, PYs from different manufacturing processes were well correlated (*r* > 0.74); very rarely did templates switch between the extreme ends of high and low manufacturability across processes (< 0.5%, Fig. 2C). Therefore, we used the consensus PY calculated across all processes for downstream analysis.

To confirm that PYs derived from pooled libraries agreed with the yields generated by single-template IVTs, we performed individual IVT validation on 44 randomly selected mRNAs distributed along the PY range, using a batch process (Methods 2.2). The IVT yields generated by individual templates tracked with the PY low-medium-high groups, supporting the use of pooled, high-throughput measurements for downstream analysis (Fig. 2D).

### Drivers for mRNA manufacturability are multi-factorial and complex

Poor manufacturability may result from low IVT yields and/or high truncation rates^34–36^. To determine the impact of high truncation rates, we captured the locations of 3’ends by ligating a 3’end adapter to the IVT products and short-read sequencing after reverse transcription (Methods 5). The high-, medium-, and low-PY transcripts showed similar truncation rates of 11.24%, 12.08% and 12.58%, respectively (Fig. S4A). Albeit slightly increasing with decreasing PYs, measured truncation rates were too low to be the primary determinant of full-length IVT yield. Additional unknown sequence features may play an important role in determining high- versus low-RNA manufacturability.

To further explore potential manufacturability drivers, we tested for differential representation of 1,386 sequence features in high- *vs.* low-PY DNA templates and secondary structure features measured using SHAPE-MaP^37^ (Methods 6, Fig. S5, Table S3). Overall, 87% (1,208/1,386) of the features tested had significantly different representation between high- and low-PY templates, indicating a clear difference in primary sequence between the two groups (*p*-values <0.01 after correction for multiple testing, Table S4). Low-PY templates showed higher T content, lower sequence complexity, and higher presence of repetitive sequences compared with high-PY templates. Secondary structure showed a linear trend of increased shape reactivity (corresponding to RNA structuredness) between low and high PYs, indicating greater structure with lower PY. Absolute differences in mean shape reactivity were highly significant (p-values <10^−50^) but small, indicating that the abundance of secondary structures negatively impacted the efficiency of transcription by T7 polymerase (Fig. S5A), but is not sufficient to explain differences in manufacturability.

To summarize RNA manufacturability drivers, we used LASSO regression to rank features by their explanatory power. The best correlation between predicted and measured PYs was *r* = 0.51 and required 900 of the original 1,386 features (Fig. S5B, Fig. S5C, Table S5). This low predictive power, the high number of selected features and their complex interrelationships motivated us to develop an interpretable deep-learning model to predict PY directly from the template’s primary sequence.

### A MAP-Net deep-learning model accurately predicts pseudoyields and identifies sequence motifs that reduce manufacturability

To enable PY predictions from a DNA template, we developed a Multi-Scale Attention Projection Network (MAP-Net) model trained on PY values (Fig. 3A, Methods 7). An attention layer in the model’s architecture allows it to identify relevant regions of the sequence for PY prediction, making the model interpretable^38–39^. We used 80%, 10% and 10% of the templates to train, validate, and test the model, respectively, with 10-fold cross-validation. The correlation between predicted and measured consensus PYs (*r* = 0.78 ± 0.01 Fig. 3B) was higher than amongst PYs measured from different IVT processes, indicating good accuracy. The model accounted for 61% of data variance, with a symmetric mean absolute percentage error (SMAPE) of 12% (21%, 9% and 17% in the low-, medium- and high-yield classes, respectively). Percentage errors of the model compared to average coefficients of variation in the measured data (see Fig. 2A) suggested that most of the variance not explained by the model should be attributed to noise in the experimental data. For predicted PY values greater than 1.10/1.25, less than 10% /1% of the templates show low measured PYs, respectively (Fig. 3C). These results suggest that MAP-Net can be used reliably to identify problematic sequences for the IVT process.

**Figure 3:**
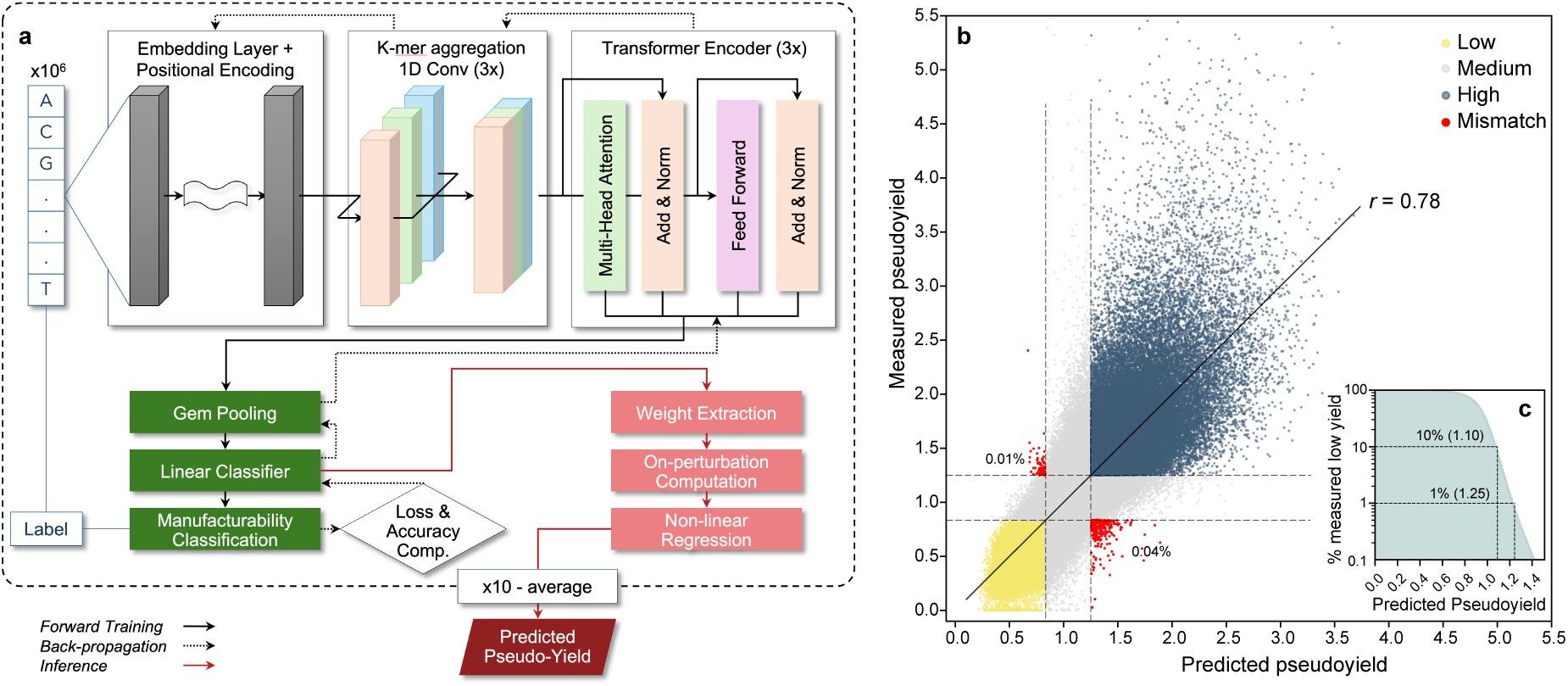
Multi-Scale Attention Projection Network (MAP-Net). **A.** MAP-Net takes nucleotide sequences as input and uses a combination of 1D Convolutional layers and Transformer encoders to learn the embeddings of the sequences. MAP-Net is trained as a binary classifier, with an 80%-10%-10% training-validation-test split of the dataset, using as a label the consensus PY across IVT processes dichotomized on the median of the PY distribution. After training, the final layer embeddings are projected onto the normal vector of the decision boundary to generate a continuous-valued predicted PY for new sequences. The process is repeated 10 times removing each time from the training-validation set a different 10% of the data, and the 10 predicted PYs are averaged to generate the output value. **B.** Main: scatterplot of predicted *vs.* measured PYs for templates in the test sets of each of the 10 trainings performed on a different train-validation set, covering the entire template library. Templates with concordant predicted and measured high/low PYs are shown in blue/yellow, respectively. Templates with predicted and/or measured medium PY are gray. Templates with discordant predicted and measured low/high PYs are shown in red. The solid line is a linear regression of predicted *vs.* measured PYs. **C.** Inset: percentage of templates in the library with measured PY in the low range (<0.834), for predicted PY values greater or equal than a given threshold.

No template in the training set included any of the most common T7 terminator sequences (Table S6, all search e-values >1) known to cause the T7 RNA polymerase enzyme to pause and/or fall off the DNA template during IVT^23^. To verify MAP-Net’s ability to learn general IVT molecular mechanisms, we introduced six T7 terminator sequences^40^ with diverse, experimentally determined termination efficiencies into 45 randomly selected high-PY templates, for a total of 270 modified templates (Fig. 4A). The PY reduction predicted by MAP-Net correlated well with the experimental data (*r* = 0.92, Fig. 4B). Despite not having been trained on any of those sequences, MAP-Net recognized accurately the added T7 terminator sequences as responsible for decreased PYs, as highlighted by the attention values along the respective templates (Fig. 4C).

**Figure 4:**
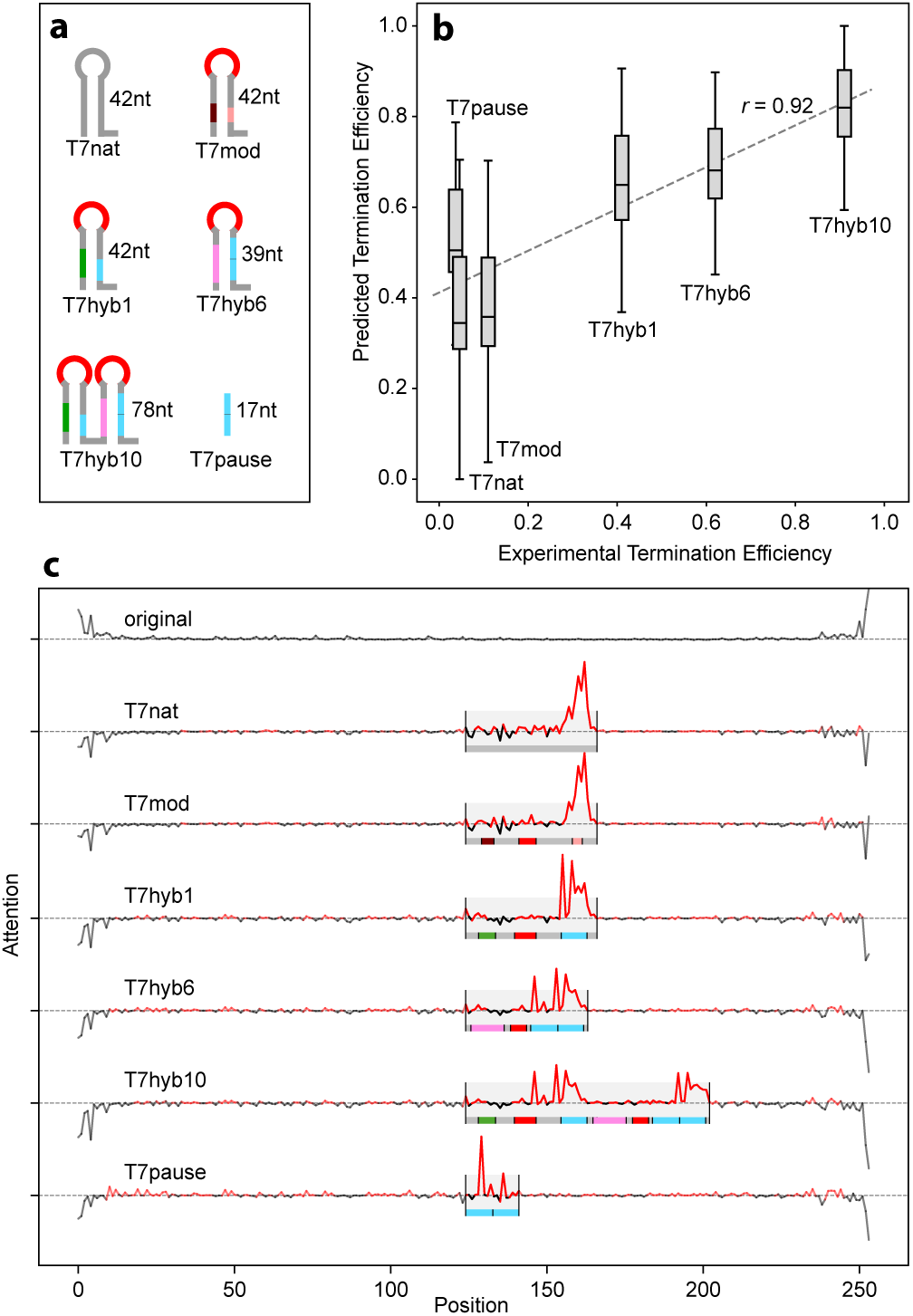
MAP-Net prediction of T7 terminator efficiencies. **A.** Diagrams of 6 known T7 terminator sequences^30^ that were each individually inserted into 45 randomly selected, high-PY templates, starting from the same template position (nucleotide 120). Numbers show the length of each terminator sequence in nucleotides. Similar colors indicate similar sub-sequences. **B.** Box-plots for the termination efficiency predicted by MAP-Net for the 45×6 = 270 templates, defined as predicted PY for the template with the inserted terminator divided by the predicted PY for the original template, are shown *vs.* the termination efficiency experimentally determined for the respective terminator^30^. Boxes represent medians and interquartile ranges (IQRs), whiskers are 1.5*IQR. The dashed line is a linear regression of predicted *vs.* measured termination efficiencies. **C.** Example of a typical test sequence. The values produced from MAP-Net’s attention layer, proportional to the relevance attributed by the network to each nucleotide for the low-PY prediction, are shown on the y-axes *vs.* the nucleotide positions in the DNA template. The first row shows the attention of the original sequence; subsequent rows show the difference between the attention of each template modified with the insertion of a terminator, and the original template. Red/black indicate positive/negative attention, *i.e.* sequences responsible for low-/high-PY, respectively. Shaded areas indicate the region in which each terminator was inserted; color coding for sub-sequence portions matches panel (a).

To search for additional T7 terminator sequence motifs, we performed motif analysis on all termination events observed in the RNA 3’end capture sequencing dataset described above. Most contained the 7-nucleotide motif ATCTGTT previously associated with class II T7 terminators, or variants thereof (Table S7). To determine whether MAP-Net can also estimate PYs when T7 terminator variants are inserted into different sequences, we inserted three terminator motifs (ATCTGTTA, ATCTGTTT, TTCTGTTT) into 6 different DNA templates and validated the transcription efficiency using batch IVT. The selected motifs induced termination in a context-specific manner (Fig. S4B). Predicted PYs correlated with experimental results (*r* = 0.66, p-value < 10^−3^, Fig. S4C), indicating that MAP-Net predicted the effect of class II terminators and their variants in a context-dependent manner.

### A MAP-Net based genetic algorithm tunes manufacturability over >7.5-fold range for two mRNA therapeutic products

To test the usability of our AI model for real-world mRNA drug products, we developed a genetic algorithm (GA), which uses MAP-Net to mutate a DNA template at synonymous positions to increase or reduce the predicted PY (Fig. S6A, Methods 8). We applied the GA to XBB.8 Spike protein, a recent variant of SARS-CoV-2 (3.8 kb) widely used as antigen in approved COVID-19 vaccines^41^ and hSpCas9 (human codon-optimized Streptococcus pyogenes Cas9, 4.3 kb) a construct used in *in vivo* mRNA-CRISPR-Cas9 gene editing therapies and *ex vivo* cell editing for mRNA-CAR-T applications^13^. Starting from the original sequences, we used the GA to design products with different predicted PYs, including two RNAs with poorer manufacturability and two RNAs with better manufacturability. The GA introduced a variable number of mutations to steer the predicted PY as directed (Fig. S6B). We then added 5’ and 3’UTRs to the sequences and performed batch IVT. We observed a very high correlation between predicted and measured IVT yields for full-length products (*r* ≥ 0.98, Fig. 5A), indicating that MAP-Net can be used to optimize IVT yields for real-life mRNA therapeutics. When starting from a naturally occurring sequence, the Spike XBB.8 gene, the GA reduced full-length yield up to 74% or increased it up to 101%, for a 7.6-fold yield range. When starting from an already highly optimized sequence, the hSpCas9 gene, the algorithm could reduce full-length yield by 85%, but increased it by 18%, for a 7.8-fold yield range.

**Figure 5:**
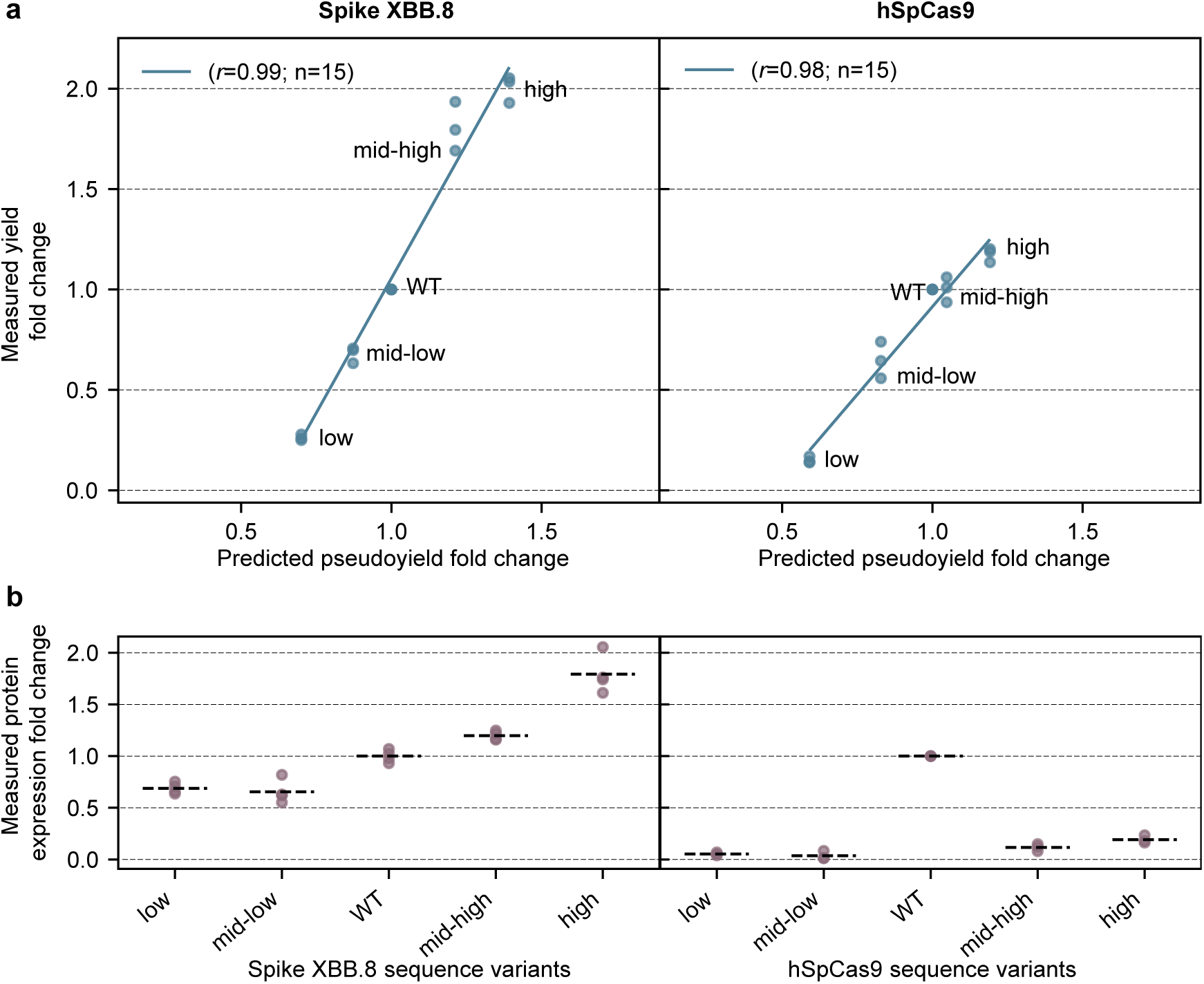
MAP-Net based manufacturability tuning of two mRNA drug products. A Map-Net-based Genetic Algorithm was used to produce synonymous sequence variants with different predicted PYs (low, mid-low, mid-high, high) for two mRNA products: the Spike protein from the SARS-CoV-2 XBB.8 strain (left) and the hSpCas9 gene construct (right). Original sequences are indicated as Wild Type (WT). **A.** For each sequence variant, the yields of full-length IVT products are shown *vs.* the predicted PY. Both values are reported as a fold change *vs.* the WT sequence. Three replicates per sequence variant are shown as dots, the solid line is a linear regression of measured *vs.* predicted data. **B.** Fold changes of protein expression measured upon transfection of 293T cells with the various sequence variants produced using 100% m1Ψ. The dotted line represents the average value between replicates.

To assess the impact of our sequence optimization procedure on the translational efficiency of mRNAs inside cells, we repeated the IVT processes using 100% N¹-methylpseudouridine (m1Ψ, Methods 9). The RNA products, with m1Ψ, 5’ and 3’UTRs, 5’ cap and poly(A) tail, were transfected into 293T cells using messenger-max lipofectamine, and protein expression levels were determined (Fig. 5B). For Spike XBB.8, lower yield constructs also had lower intracellular translation, consistent with the presence of truncated mRNAs, while higher yield constructs showed up to a 69% increase in protein production. For hSpCas9, all modified constructs had significantly lower intracellular translation, indicating a negative impact of the GA on sequence motifs that optimize translation efficiency. However, in both cases the amount of protein produced was positively correlated with predicted yield, suggesting the possibility to co-optimize concurrently for manufacturability and translation (Fig. S6C).

### Human transcripts have medium-low pseudoyields, and transcription initiation rate could be a main driver for manufacturability

To determine what percentage of natural transcripts would benefit from codon optimization to improve manufacturability, we used MAP-Net to predict the PYs of 19761 primary human transcripts in the ENSEMBL database (Methods 10). Human transcripts had median PY of 1, 2,5% (501 transcripts) had predicted PY <0.825, *i.e.* low manufacturability, and <0.03% (only 4 transcripts) had a predicted PY greater than 1.225 (high manufacturability). This suggests that the majority of the human transcriptome would benefit from codon optimization to develop better RNA medicines. Interestingly, the 100 transcripts with the lowest predicted PYs showed a strong enrichment for genes associated with the sensory perception of smell (*p*<10^−5^, Figure 6a). These genes have been reported to have high AU content and short UTRs^42^, consistent with our identification of AU content as a leading feature associated with low manufacturability (Supp. Figure 5), highlighting that manufacturing of this group of transcripts might particularly benefit from codon optimization.

**Figure 6.**
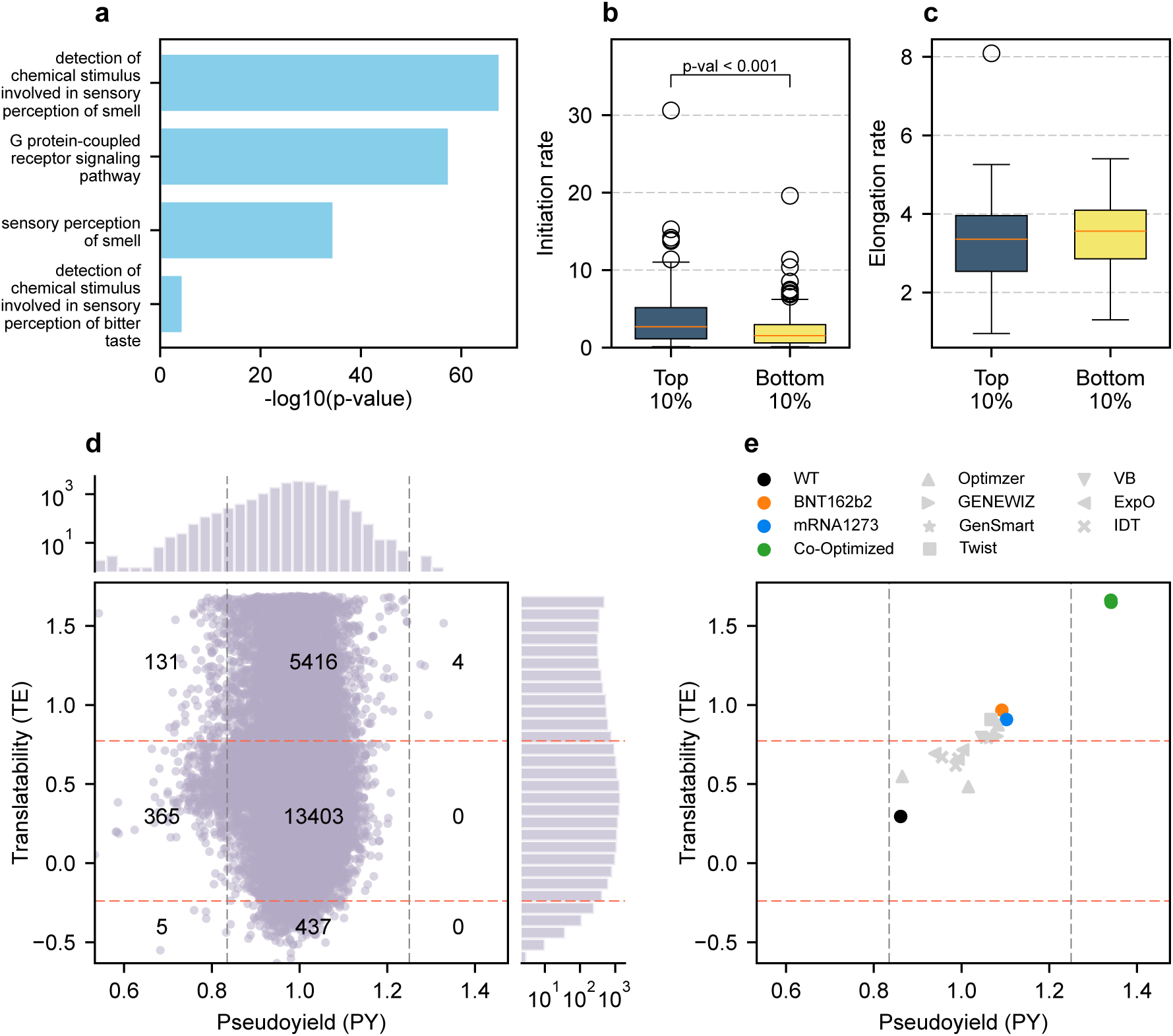
MAP-Net can be used to co-optimize manufacturability and translatability to generate better drug products. **A**: GO term enrichment of the 100 primary transcripts in the human transcriptome with the lowest manufacturability score; shown are the 4 terms with the most significant p-values. **B, C**: distribution of transcription initiation rates (b) or transcription elongation rates (c) for primary human transcripts with top or bottom 10% predicted PY. **D**: Predicted PY and TE in the human transcriptome for 19761 primary transcripts from the ENSEMBL database; dotted lines delimit low-medium-high manufacturability (gray) and translatability (orange); numbers in each cell indicate the corresponding number of human transcripts. **E**: Predicted PY and TE for different codon optimizations of the Spike protein CDS from the SARS-CoV-2 Wuhan-Hu-1strain; wild type sequence (black), codon-optimized sequences using seven commercial software products (grey, see Methods 10.4), BioNTech (orange) and Moderna (blue) commercial vaccine sequences, co-optimized sequences using a genetic algorithm based on MAP-Net and RiboNN (green).

While our predicted PY are derived from T7 polymerase in vitro transcription yields, we explored the relationship between PY and transcription initiation and elongation rates in human cells. Contrasting published transcription initiation and elongation rates^43^ between the human transcripts with top and bottom 10% predicted PYs, we did not observe a significantly different elongation rate (Bonferroni corrected p-value > 0.06), but transcripts with high PYs showed a significantly higher transcription initiation rate as compared to transcripts with low PYs (Bonferroni corrected p-value <0.001, Figure 6b,c). As transcript stoppages only account for 12.58% of low PYs (Fig. S4A), we hypothesize that transcription initiation could be a main driver for manufacturability in our dataset. Despite the extensive amount of regulation in human PolII transcription, this data suggests that similar sequence composition could impact both PolII and T7 polymerase initiation during transcription.

### Co-optimization for manufacturability and translation efficiency improves over state-of-the-art commercial mRNA products

We had previously observed that optimizing for increased manufacturability on natural transcripts can result in increased protein production, but also that manufacturability optimization on sequences already optimized for translatability can dramatically reduce it (Figure 5b, left and right panels, respectively). To further explore the relationship between PY and translation efficiency inside cells, we used the RiboNN algorithm,^44^ to predict translation efficiency (TE) for the human transcriptome (Methods 10). RiboNN was previously trained on large amounts of ribosome profiling data in different cell types and, as we did for manufacturability, we defined “low-”/“high-translatable” the first/last sextile of the training distribution, respectively. Interestingly, we observed no meaningful correlation between PY and TE in natural human transcripts (*r* = 0.06, Figure 6d). However, in contrast with what we observed for manufacturability, 28% of the human transcripts showed high translatability, suggesting that the selection pressures on manufacturability and translatability are likely to be different on natural transcripts, and that we could potentially co-optimize manufacturability and translatability on RNA sequences.

To test whether we can effectively co-optimize a real-world medicinal product, we compared manufacturability and translatability from different codon optimizations of the same Spike sequence from the SARS-CoV-2 Wuhan strain. We predicted PY and TE from the wildtype (WT) sequence (PY=0.86, TE=0.30), from sequences optimized using seven different commercial software programs, from the Moderna (mRNA 1273) and BioNtech (BNT162b2) commercial SARS-CoV-2 vaccines, and from our GA that co-optimized the WT sequence using simultaneously MAP-NET and RiboNN (Figure 6e). All codon optimization software programs improved both manufacturability and translatability of the WT Spike sequence, but resulted lower than BNT162b2 and mRNA1273 which had the highest PY (1.09, 1.10, medium manufacturability) and TE (0.97, 0.91, high translatability). By co-optimizing the WT strain with MAP-Net and RiboNN, we obtained the only sequences showing both high manufacturability and high translatability (PY= 1.34, TE=1,65), more than doubling the increase of the commercial products *vs.* the WT. Results indicate that intracellular translation and IVT manufacturability can be co-tuned for optimal drug development, significantly improving over the current state of the art.

## Discussion and Interpretation

We identified design features associated with high mRNA manufacturing yield and integrity, complementing current sequence design focused on improvements in intracellular protein production or reducing immunogenicity^43–47^. We then trained an AI model that predicts mRNA manufacturability and enables codon optimization during sequence design to maximize yield and quality independent of the manufacturing process. Our framework decouples mRNA product design from manufacturing specification, unlocking the versatility of the mRNA platform ^45^.

Pseudoyield, a yield and quality measure derived from direct Nanopore sequencing data, revealed a complex picture. The yield of full-length IVT-RNAs showed a broad, >100-fold range, indicating that product design can have a major impact on manufacturing. The lowest and highest sextile in the distribution, termed “low-” and “high-manufacturability” templates, revealed a marked difference in sequence composition. While certain differentiating features confirmed expectations, with low manufacturability sequences associated with more abundant secondary structures, T-rich repetitive elements and lower sequence complexity, a manageable number of features was not sufficient to account for the observed PY differences. Known terminators of T7 RNA polymerase accounted for only ∼10% of the poor yields. Additionally, a regression model failed to predict template manufacturability with acceptable accuracy, and the set of non-redundant sequence features was too large to provide guidelines for template design.

The observed complexity of PY results led us to develop and train an interpretable deep learning model, MAP-Net, which could predict PYs out-of-sample with over 78% Pearson’s correlation across the whole experimental range. The aggregation of multiple k-mer sizes in the encoding layer of a relatively small network, with less than 1 million parameters, was critical to achieve this precision, suggesting that sequence features at different scales play an important role in IVT. MAP-Net accurately predicted, in diverse sequence contexts, the effect of 6 control sequences known to negatively affect IVT^37^. The values of the network’s attention layer confirmed that the model recognised exactly the sequence motifs responsible for reduced manufacturability, despite not having been trained on any of the control sequences.

MAP-Net also predicted with high precision the manufacturability of two long, real-world mRNA products (∼4 kb) whose sequences had been synonymously mutated by a genetic algorithm. This indicates that the information captured in the network embeddings, even if trained on ∼20 times shorter sequences, is transferable to long mRNAs. Therefore, MAP-Net’s attention values across low-manufacturability templates can provide a useful reference to further explore sequence motifs impacting mRNA manufacturability. Applying MAP-Net on the human transcriptome revealed that most human transcripts would benefit from codon optimization to increase manufacturability, indicating that manufacturability is probably not a feature that nature transcripts are under selection pressure for. Extremely poorly manufacturable transcripts are enriched for olfactory sensing genes, which show unique sequence features including high AU content, which is one of the top features that predict low manufacturability in our data.

Since COVID-19, alternative manufacturing approaches have been investigated to increase efficiency, flexibility, and scalability for mRNA manufacturing^40^. We used diverse IVT processes and derived a multi-process consensus to identify universal features that enable RNA manufacturability for established and evolving processes. Extreme disagreements among processes were rare, with <0.5% templates having low-manufacturability in one process and high in another. At intermediate PY ranges, variations were greater, suggesting that training focused on a single IVT approach could increase MAP-Net’s precision.

Our PYs should not be used to compare the efficiency of different IVT processes. We started library preparation from each process with equal amounts of IVT-RNAs, which were not proportional to yields for each process. We also applied the same post-IVT column purification protocol to all IVT-RNA pools, ignoring individual manufacturer-specific purification strategies. Future work, using fully purified RNAs from each manufacturing process, may help better characterize and compare IVT processes. Within the limitations of our data, though, results diverged progressively from batch IVT as processes moved towards a continuous flow.

While manufacturing high-yield and high-quality mRNAs is important, the mRNA must be translated efficiently inside cells. Using a naturally occurring sequence, the MAP-Net-based codon optimization tuned both manufacturability and translatability inside cells, indicating that designing for manufacturability can also enable high translatability. By further co-optimizing for both manufacturability and translatability for the SARS-CoV-2 SPIKE sequence, we show that we can design the sequence to have even greater manufacturability and translatability as compared to existing drug products, highlighting the potential to couple manufacturability with protein production broadly for better drug design.

Fundamental modifications in the IVT process, such as the use of modified nucleotides and/or optimized polymerases, may necessitate future MAP-Net optimization via re-training or transfer learning. Still, the results presented here lay the groundwork for democratizing access to mRNA product design, in a manner similar to how very large scale integration rules enabled the current state of semiconductor-based product design. Removing the need for expensive and complex product-specific technology transfers should reduce risk and help pave the way for an RNA therapeutics revolution by accelerating the development of RNA-based medicines for all those in need.

## Supporting information

All Supplemental Table Legends, Supplemental Tables S2, S3, S5

## Acknowledgments

Miles Benton and Scott Hickey provided access to ONT technology. Kiat Yee Tan, Zhang Yu, Liya Chiu, Aurora Odierno and Edoardo Renato Bucalo contributed to study conduct. Francois Hallac conducted flow IVT. Marc Penders oversaw semi-continuous IVT. Adithya Nair, Kate A. Loveday, Emma N. Welbourne and Caroline Evans conducted batch mRNA manufacturing and analysis. Mabrouka Maamra coordinated and oversaw batch mRNA activities. Amin Abek and Purnima Mala supported sample preparation. Lisa DeTora provided writing and editorial support.

## Funding

This work was supported by Wellcome Leap as part of the R3 program.

## Author contributions

Conceptualization: DM, YW, KCL, GC, GM, RH, TvdH

Methodology: DM, YW, RH, GM, GC, CM, PFdS

Investigation: KCL, GM, GC, RH, CYT, NTXO, TYS, TWL, CM, MD, ZK, CMak, AvA,

Visualization and Interpretation: KCL, GM, GC, RH, TvdH, CYT, NTXO, PFdS, TYS, TWL, MD, AvA, YW, DM

Funding acquisition: DM, YW

Project administration & Supervision: DM, YW, PFdS, ZK, CMak, AvA, RH,

Writing – original draft: GM, GC, KCL, YW, DM

Writing – review & editing: All authors

## Competing interests

PFdS, NO, and CT are employed by Oxford Nanopore Technologies; CM is co-founder and CSO of Waterfall Scientific; CM is co-founder of Centillion Technology Ltd; SvA owns RiboPro, an RNA technologies company and Mercurna, an RNA drug development company; during study conduct, DM was an employee of Wellcome Leap. DM, GC and GM are employees of Toscana Life Sciences. DM is also employed by Pitt BioForge.

## Data and materials availability

Any methods, additional references, supplementary Figures and tables, details of author contributions and competing interests, and statements of data and code availability are available at https://doi.org/XXX.

## Additional Materials provided

Materials and Methods

Extended Data: Figures S1 to S6

Supplementary Tables S1 to S7

## Materials and Methods

### 1. Library of 1 million diverse DNA templates

A library of 1 million DNA sequences was designed to include both coding and non-coding sequence fragments derived from multiple organisms, spanning multiple kingdoms such as viruses, bacteria, mammals, birds and fungi, as illustrated in Fig. S1A. Multiple genome assemblies were fragmented using a moving window of 254bp and a stride of 10bp, and templates with identity > 90% were removed using CDHIT^48^. From the remaining fragments, 1 million sequences were selected to create the final library.

For each fragment, a 26-nucleotide long T7 polymerase promoter (TAATACGACTCACTATAGGGAAATAA) was added at the 5’ end, and a 20-nucleotides poly-A tail was added at the 3’ end, resulting in a library of 1 million templates, each 300 nucleotides long (Fig. S1B). As an internal control, 13 natural and synthetic T7 polymerase terminator sequences, which have been previously described^40^, were introduced at position 150 of 300 randomly chosen templates. This resulted in a total of 3,900 sequences containing known T7 polymerase terminators. The final library contains highly variable fragments that cover 100% of the combinatorially possible 11-mers and 95% of the possible 12-mers (Fig. S1C).

### 2. DNA template preparation and in-vitro transcription (IVT)

#### 2.1 Pooled IVT

All DNA templates included in the library just described were purchased from Twist Bioscience as single stranded DNA oligo pools (300bp) and PCR-amplified using RM 1F and RM 1R primers (Supplementary Table 1). Amplified DNA templates were then to be gel extracted and used as input for four IVT protocols: batch using a commercial kit, batch using in-house materials, semi-continuous and continuous. In all pooled IVT protocols, IVT-mRNA pooled products were purified using NEB Monarch RNA purification columns.

##### 2.1.1 Pooled Batch IVT using a commercial kit

The NEB HiScribe kit was used according to manufacturer’s instructions at GIS. In brief, for 300bp transcripts, 10ng DNA templates were mixed with 7.5mM NTPs and 0.5uL T7 RNA polymerase mix in 20uL (final concentration is 0.75X reaction buffer) for 4h at 37°C.

##### 2.1.2 Pooled Batch IVT using an optimised in-house composition

Small-scale IVT reactions were performed by University of Sheffield, UK, at a final concentration of 0.00193 µg/µL in a 20 µL reaction. The reaction contained 10 mM each of ATP, CTP, GTP, and UTP (Roche), 42 mM magnesium acetate, 40 mM HEPES, 2 mM spermidine, 10 mM dithiothreitol (DTT), 50 mM sodium chloride (NaCl), and 0.001% Tween 20 in an in-house-made IVT reaction buffer. T7 RNA polymerase (Roche), inorganic pyrophosphatase (Roche), and RNase inhibitor (Roche) were added to the reaction mixture. All reactions were assembled on ice and incubated at 37 °C for 4 hours.

##### 2.1.3 Pooled Semi-Continuous IVT

Small-scale IVT reactions were performed at RIBOPRO (Oss, The Netherlands) with in-house developed and produced RiboPerfect™ RNA Polymerase, which is a novel enzyme with improved RNA product yield and with a reduction in byproducts, such as double stranded RNA. A proprietary IVT reaction buffer was used with a final DNA concentration of 15 ng/µL, 10 mM of ATP and 7.5 mM of CTP, GTP, and UTP (Thermo Fisher Scientific). Inorganic pyrophosphatase (Thermo Fisher Scientific), and RNase inhibitor (Thermo Fisher Scientific) were added to the reaction mixture to 0.005 U/µL and 1U/µl final concentration, respectively. All reactions were assembled at room temperature and subsequently incubated at 37 °C for 3h and afterwards diluted to ∼1 µg/µl.

##### 2.1.4 Pooled Continuous IVT

Continuous IVT in flow conditions was achieved using a patented process (Centillon, Cambridge, UK)^49^. The setup comprises two independent feed streams that continuously feed into a microfluidic bioreactor where the output was continuously collected until the required quantity of RNA is generated. The flow rates of the two independent feed streams, which contain the reagents required for efficient mRNA synthesis, were set to maximise the yield and purity of the final drug substance. The net flow rate is calculated based on the reactor volume and residence time. For a reactor volume of 3.6 mL, the net flow rate ranges from 30 to 120 µL/min for a residence time of 30 minutes to 2 hours. To reduce the potential for degradation by hydrolysis to double-stranded (ds) RNA, the RNA was collected as soon as it is formed within the channels of the flow bioreactor at the end of the residence time.

#### 2.2 Individual IVTs

For individual gene fragment IVT validation (see Fig. 2D), double-stranded DNA fragments were synthesized and amplified using RM 1F and RM 1R primers. Subsequently, gel extraction was performed to purify DNA templates at the correct size. Then 20ng DNA templates were used for batch IVT using a commercial kit (NEB HiScribe), as described in 2.1.1. After 4h incubation at 37°C, DNase treatment was performed and IVT products were purified by NEB Monarch RNA purification columns.

For SARS-CoV2 XBB.8 Spike (K982P and V983P were introduced to stabilize spike as pre-fusion state) and hSpCas9 validation experiments (see Fig.5 and S6), full-length constructs were synthesized and IVT DNA templates prepared by gel extraction of amplified PCR products using RM 2F and RM 2R primers. Next, batch IVT was performed using 50ng DNA templates mixed with 10mM NTPs and 2uL T7 RNA polymerase mix in 20uL (final concentration is 1X reaction buffer) at 37°C for 2h. After DNase treatment, all these IVT reactions were purified by NEB Monarch RNA clean-up columns and RNA concentrations were measured by NanoDrop spectrophotometer or Qubit Fluorometers. To estimate the concentrations of full-length transcript (adjusted concentrations), concentrations obtained from NanoDrop spectrophotometer or Qubit Fluorometers are multiplied by the percentage of full-length transcript. This percentage is calculated by dividing the signal of full-length transcript by the sum of signal from full-length transcript and truncated IVT products. These signals are determined by quantifying band intensity in gel electrophoresis results by densitometry analysis. Alternatively, the relative abundance of full-length transcript versus truncated IVT product are assessed by fragment analyzer. All primers and sequences are provided in Table S1.

### 3. Library preparation and next-generation sequencing

#### 3.1 Long-read sequencing

Both DNA template library and IVT product were deep sequenced using Oxford Nanopore Technologies. Ligation sequencing Kit V14 (SQK-LSK114) and Direct RNA sequencing Kit (SQK-RNA004) were used to prepare DNA and RNA libraries respectively. Subsequently, the DNA library was sequenced using FLO-PRO114M (R10.4.1) flowcells in triplicate to ensure reproducibility and robust genomic coverage. RNA libraries resulting from each IVT manufacturing process were sequenced in duplicate using FLO-PRO004RA flowcells. All sequencing runs were performed with live basecalling on a PromethION sequencing instrument using the <u>dna_r10.4.1_e8.2_400bps_sup@v4.2.0</u> model for DNA libraries and the <u>rna004_130bps_sup@v3.0.1</u> model for RNA libraries. The resulting runs are listed in Table S2. Raw data is available for download at [link to be provided].

#### 3.2 Short-read sequencing

To prepare libraries for Illumina sequencing in order to determine transcription stoppage sites, a modified version of NEB small RNA library preparation (E7330) protocol was adopted. In brief, purified IVT products were first ligated with the 3’ adaptor and subsequently ligated with 5’ adaptor. Then RT primer was annealed for reverse-transcription and PCR amplification was performed. Amplified libraries were subjected to TBE-polyacrylamide gel electrophoresis and size-selection was performed to extract 150-500bp region. Purified libraries were submitted for paired-end 150bp Illumina sequencing.

### 4. Data processing

#### 4.1 Sequencing data filtering and quantification

Raw Oxford Nanopore sequencing reads were processed using **Minimap2**^50^ and **Samtools**^51^ to generate sorted alignment files. Minimap2 was used for read mapping with the ***-ax map-ont*** preset, optimized for Oxford Nanopore long reads. Reads were mapped against the reference library using the one million template dataset, and the resulting alignments were sorted using **Samtools sort command**. To ensure that only complete fragments were counted for DNA and RNA quantification, aligned reads were filtered based on a minimum coverage threshold for the input templates. For DNA data analysis, reads that did not cover the interval between position 10 and 280 were discarded. For RNA data, only reads covering the interval between position 40 and 280 were retained. These ranges account for variability in length due to the poly-A tail at the 3’ end and transcription initiation after the T7 promoter in the RNA data (position 26+13). After filtering, the raw counts were normalized by the total number of reads for each replicate and converted to counts per million (CPM).

#### 4.2 Pseudoyield (PY) calculation

DNA quantification data was filtered to remove templates where the calculated CPM was zero or close to zero, ensuring that only templates with sufficient read depth were retained. To determine the appropriate CPM cutoff threshold for reliable quantification, the three DNA replicates were compared. A range of thresholds, from 0 CPM to 1 CPM, was applied, and the percentage of shared templates below each threshold was calculated. This analysis helped identify the optimal tradeoff between retaining shared templates and discarding low-confidence ones (Fig. S2C). As a result, a threshold of 0.2 CPM was applied to retain those templates where all three replicates had CPM values above the threshold, retaining 780,832 templates (Fig. S2D). As a result of the filtering process, the coefficient of variation (CV) median value for the retained templates along replicates decreased from ∼10 to ∼7 (Fig. S2F). No additional filtering was applied to RNA data. DNA and RNA CPM were then used to calculate the PY as a measure of the number of full-length RNA molecules that are transcribed by the T7 polymerase from a single DNA molecule.

The three DNA and eight RNA quantification as CPM were used to calculate the PY and its variance for each individual template using a second order Taylor expansion (Equations 1 and 2, respectively^52^:

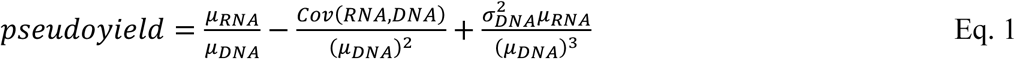

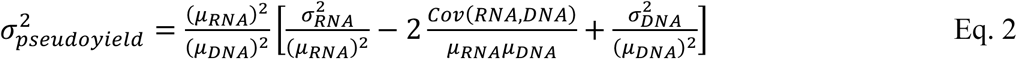

Where:

- 𝜇*_DNA_*: Mean DNA CPM across replicates
- 𝜇*_RNA_*: Mean RNA CPM across replicates
- 𝐶𝑜𝑣(𝑅𝑁𝐴, 𝐷𝑁𝐴): Covariance between RNA and DNA CPM across replicates
- 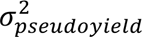: Variance of PY
- 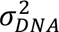: Variance of DNA CPM
- 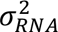: Variance of RNA CPM.

This PY (Eq. 1) was used instead of a simple 𝜇_&’(_/𝜇_*’(_ ratio to correct for bias, due to noise and uncertainty in quantification. By accounting for the covariance between RNA and DNA the shared noise is removed, and by accounting for DNA variance, error propagation from denominator variability is reduced.

The PY variance (Eq. 2) was estimate using second-order error propagation (Taylor expansion). This quantifies how variability in RNA and DNA replicates, as well as their covariance, contribute to the overall uncertainty in the PY estimate.

#### 4.3 Correlation analyses

All pairwise correlation analyses in this manuscript were Pearson correlations, calculated using the *DataFrame.corr()* method in pandas (v2.2.3) and reported as Pearson’s correlation coefficient “*r*”. Each correlation was computed consistently with this approach to ensure comparability across datasets.

### 5. Analysis of IVT-mRNA Termination through short-read sequencing

#### 5.1 Identification of premature termination

We aligned the sequencing results of the individual IVT runs with the reference library. To identify early terminations of IVT, we ascertain the presence of reads containing a 3’ sequencing adapter at a site other than the 3’ end of the reference sequence. As no fragmentation of the IVT product was performed, such early 3’ adapter fusion sites are indicative of premature IVT termination.

This inventory of premature termination sites allowed us to build a comprehensive map of sequence motifs associated with early IVT termination. Our analysis focused on all 4096 possible 6-mers and their association with early termination. We pursued two independent approaches to identify which motifs induce IVT termination and should be avoided in mRNA sequence design. Both approaches are variants of meta-gene analysis.

#### 5.2 Motif occurrence relative to termination events

In our first approach, we created an alignment of all termination sites. We then analyzed the resulting alignment for occurrence of any 6-mer motif at positions up to 50 nt preceding the termination site. If a motif shows weak or no association with termination, it will occur randomly with respect to the termination site and the resulting distributions of motif positions will appear generally uniform. For motifs showing strong association with termination, they will be positioned preferentially at a certain distance from the termination site and thus the distributions will show one or more distinct peaks. We used Shannon entropy (S) to quantify the uniformity of these probability distributions p(x) over N discrete positions x.

The Shannon entropy of a discrete probability distribution is given as:

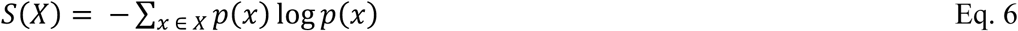

And thus, a uniform distribution over N discrete bins will have a maximal entropy value of:

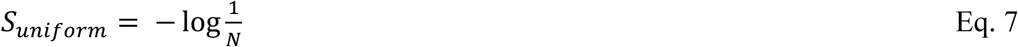

We can define a measure of excess entropy as a distance from this uniform case:

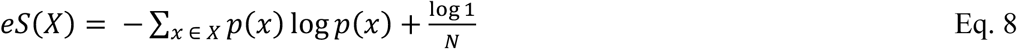

This excess entropy eS is thus 0 in the case of uniform distributions and higher the further a particular distribution deviates from a uniform distribution, which allows us to quantify the strength of association of individual motifs with IVT termination events.

#### 5.3 Consequences of motif occurrence

Alternatively, we can observe the result of individual motif occurrence on the quantity of product following the motif. For every individual 6-mer motif, we plotted the remaining normalized quantity of product at each position following this motif.

If an individual motif has no association with termination, we expect constant-rate termination (e.g. from the presence of other motifs or random events) after the occurrence, which can be understood as a Poisson process and described by an exponential distribution with the parameter λ describing the rate of decay:

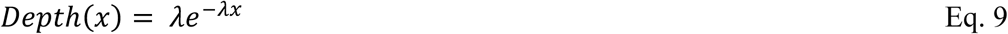

If the motif is causative for termination, we expect an additional sigmoid behavior that reduces remaining depth over a narrow range of positions:

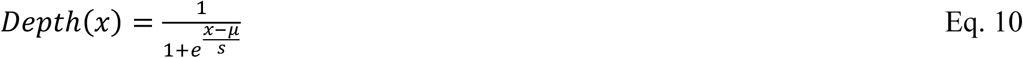

With the parameter μ describing the relative location of the termination event and parameter s describing the width over which termination occurs relative to the position of the motif.

Superimposing these processes allows us to create a mixture model that describes the drop-off of depth following an individual motif:

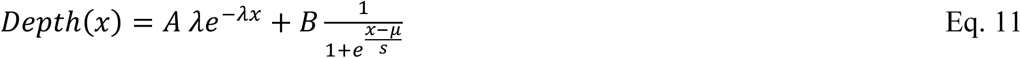

with additional parameters A and B describing the relative contributionsof the respective stochastic (A) and sigmoid (B) behaviors. This model was fitted to all observed decay curves following the first occurrence of each of the 4096 6-mer motifs. Purely stochastic termination would result in a large A parameter and a small B parameter, whereas strong association of IVT termination with a motif would result in a larger B parameter. We can thus define the termination strength of an individual motif as the ratio of B/A, which we report for numerical convenience as parts per thousand.

#### 5.4 Assessing the impact of premature termination on overall yield

To assess whether low observed PYs are caused by an overabundance of premature IVT terminations, we grouped templates according to their PY into ‘low’ (below −1σ) ‘medium’ (−1σ to +1σ) and ‘high’ (> +1σ) PY classes. We then calculated the number of transcripts terminating early (stop before the poly-A tail of the template, at or before position 280) as a fraction of total observed transcripts. We find that most templates produce high fractions of full-length transcripts, and we see a slight overabundance of early IVT termination in lower-yield classes. However, the absolute differences are small and do not convincingly explain the difference in PYs.

### 6. Analysis of features differentiating low- vs. high-pseudoyield DNA templates

#### 6.1 Primary sequence features

To define which physicochemical and sequence-derived properties most strongly predict high IVT yield, we first computed a comprehensive set of DNA-encoded features for each template in our library (see Table S3 for a complete list) for a total of 1,386 features. To assess the statistical relevance of each feature, we compared the distributions of feature values between high-PY and low-PY templates using the non-parametric Mann–Whitney U test. This test evaluates the null hypothesis that the two groups originate from the same distribution. P-values were adjusted for multiple testing using the Bonferroni correction, and features with corrected p-values below 0.01 were retained, resulting in a subset of 1,230 features selected for further analysis (Table S4).

To rank features by explanatory power, we first removed mutually correlated features with Pearson correlation > 0.90, which eliminated 22 features (listed in Table S5), yielding a final set of 1,208 nonredundant features. All feature values were standardized (zero mean, unit variance) and missing entries set to zero. We then applied an L₁-penalized logistic-regression model (LASSO) implemented in scikit-learn (v.1.2.0), using five-fold cross-validation to select the regularization strength that maximized mean ROC-AUC. The dataset was split for training (80%) and testing (20%). Features whose coefficients remained nonzero under the optimal penalty were taken as predictive variables; their relative importance was assessed by the absolute magnitude of their fitted coefficients. This approach yielded a sparse, interpretable signature of the key sequence and physicochemical determinants of high *vs.* low PY classification.

#### 6.2 RNA structure probing

To probe RNA structures, SHAPE-MAP was performed as described previously with slight modifications^53–54^. In brief, DMSO or 2A3 (50mM) (2-aminopyridine-3-carboxylic acid imidazolide) were added to the IVT reactions at 37°C 10 minutes before IVT reactions were completed. After column clean-up, purified RNA was mixed with random primer for reverse transcription in the presence of MnCl_2_. Subsequently, second strand DNA was synthesized. After the ends of the DNA fragments were repaired, the Illumina adaptor was ligated, and USER enzyme treatment performed (NEB). Finally, Illumina primers were used for library amplification and submitted for next-generation sequencing.

### 7. MAP-Net design, training and validation

#### 7.1 Data Preparation and Loading

Sequences described in section 1 were tokenized. The 10bp stride of each template was incorporated as overlap between contiguous sequences. Both the nucleotide sequences and the overlap data were converted into an integer representation. These representations were then stacked to create a combined input tensor. As a result, each sample in the dataset consists of a nucleotide sequence and the corresponding overlap encoding, merged into a single array with the associated label.

#### 7.2 Model Architecture

We developed a hybrid model^38^ that combines 1D convolutions with a Transformer-based neural network as its backbone. The architecture integrates several key components to effectively capture sequence patterns and long-range dependencies.

A custom embedding module combines nucleotide token indices with additional overlap-feature indices into a single embedding. Each nucleotide-feature combination is mapped to a unique index and embedded in a shared 132-dimensional space. If overlap features are not used, the model applies a standard embedding for nucleotide tokens alone. To preserve sequential ordering information, we incorporated a positional encoding layer^39^, specifically using a standard sine-cosine-based encoding.

The model includes a k-mer aggregation module, which uses 1D convolutions to capture local sequence patterns spanning various k-mer sizes (e.g., k = 1, 5, or 11). This enhances the ability of subsequent layers to recognize sequence motifs at multiple scales.

The core of the model consists of multiple Transformer Encoder blocks. Each block applies multi-head self-attention, followed by a feedforward sublayer. Residual connections and layer normalization are included to improve training stability. A Generalized Mean (GeM) pooling layer^55^ aggregates hidden representations across the sequence dimension, and a final feedforward linear layer transforms the pooled representation into class logits for classification.

#### 7.3 Training Procedure

The model was trained by minimizing a Cross-Entropy loss function. During each iteration, batches of sequence–overlap pairs were processed through the network, generating class probabilities that are compared against the ground-truth labels. Model parameters were updated using backpropagation using a gradient-based optimizer, such as Adam, to refine the network’s weights and improve classification performance.

Schematic workflow:

A. Data Splitting: to ensure robust evaluation, we performed 10 independent splits of the dataset into training (80%), validation (10%) and test (10%) subsets. Each test set was constructed to be completely disjoint from the others, such that no sequence appeared in more than one test set.
B. Hyperparameters: our configuration uses a batch size of 512, hidden dimension of 132, 4 self-attention heads, 3 encoder layers, and a dropout rate of 0.2.
C. Training strategy: For each of the 10 independent dataset splits, the model was trained for up to 100 epochs with early stopping and learning-rate scheduling based on validation-set performance to prevent overfitting and promote convergence.

#### 7.4 Evaluation and Metrics

The final trained model was evaluated on the held-out test set. Standard classification metrics—such as accuracy, MCC, precision, and recall—were computed to assess performance. Additionally, attention weight matrices from each Transformer encoder layer can be analysed to identify key regions or motifs within nucleotide sequences that the model considers most relevant for classification.

#### 7.5 Pseudoyield prediction

To analyse the model’s decision boundary in our binary classification task, we computed projections of input features onto the normal vector of the decision boundary. The decision boundary represents the points at which the model assigns equal probability to both classes, corresponding mathematically to:

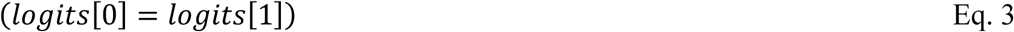

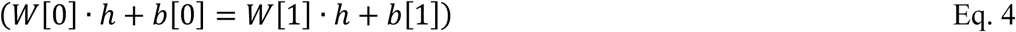

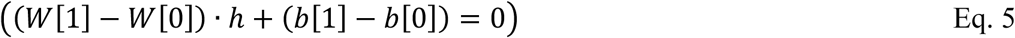

Here *logits* are the model outputs for each class, *h* is the activation vector for a given input sample, (*W*[1]-*W*[0]) defines the normal vector pointing in the direction that best separates the two classes, and (*b*[1]-*b*[0]) is the scalar offset.

The decision boundary is a hyperplane that divides the feature space into two halves, in our case *low yield* class and the *high yield* class. By projecting a sample’s activations onto the normal vector, we can effectively measure how far the sample is from the decision boundary and in which direction. A positive projection indicates that the sample lies on the side corresponding to the *high yield* class, while a negative projection indicates the *low yield* class. The magnitude of this projection reflects the classification confidence, or margin, which shows how clearly the sample is classified. We then fit a custom exponential-like function to the data by modelling the relationships between the projection values and the measured PY values. Specifically, we used non-linear fitting via SciPy’s *curve_fit* function, initializing the parameters with all zeros and setting the maximum number of function evaluations to 1000. The optimized parameters were then used to compute the predicted PY for each sequence.

### 8. Genetic Algorithm for Sequence optimization

#### 8.1 Overview

We developed a genetic algorithm (GA) to optimize nucleotide sequences with the aim of improving their yield based on the pre-trained deep learning model. The optimization process employed both evolutionary principles and advanced computational techniques to balance the performance of the generated sequences.

#### 8.2 Initial sequence and population initialization

We initialized the GA using a given nucleotide sequence (that is, the original sequence). We generated a population of candidate sequences by introducing random synonymous mutations within the allowed regions, maintaining the integrity of the amino acid sequence. Mutations were introduced randomly at the codon level, avoiding excluded regions marked as invariant, resulting in a diverse initial population that maintains functional constraints.

The fitness assessment for each candidate sequence comprised two key criteria:

Predicted PY score (function A). This metric quantifies the distance and direction of embedding, extracted from the pre-trained model, of a candidate relative to the classification decision boundary of the pre-trained transform-based neural network. Sequences with higher predicted PY scores (positive class optimization) or lower predicted PY scores (negative class optimization) indicate higher classification confidence.
Mutation minimization (C function). To avoid excessive divergence from the original sequence, we calculated a mutation-minimization score.

#### 8.3 Pareto optimization

After fitness assessment, we employed Pareto optimization to identify a subset of non-dominated individuals within the population. An individual sequence was considered non-dominated if no other sequence was superior for both fitness goals simultaneously. This strategy preserves the diversity of candidate solutions while promoting a balanced improvement between objectives.

#### 8.4 Population selection and upgrading

Elite selection was performed by directly preserving a small subset of the highest-ranking individuals according to Pareto front criteria, ensuring the preservation of the best solutions across generations. Additional individuals for the next generation were selected through tournament selection, promoting competitive but diverse breeding pools.

#### 8.5 Crossover and mutation

The GA implemented a uniform crossover technique that respects codon boundaries. Two parental sequences selected by tournament selection produced progeny through random codon exchanges at the allowed sites. Subsequent adaptive mutation introduced further variation; the mutation rate decreased linearly over generations, starting high to encourage exploration and decreasing thereafter to promote solution refinement.

#### 8.6 Metropolis-Hastings acceptance criterion

To mitigate premature convergence and explore wider solution spaces, we integrated a Metropolis-Hastings acceptance criterion. Candidates created by adaptive mutation were accepted probabilistically even if their fitness scores were slightly lower.

#### 8.7 Temperature schedule and termination criterion

The temperature followed an exponential decay schedule, promoting a wider search at the beginning of the evolution and a refinement in subsequent generations. The initial temperature was set high (e.g. 5.0) and decreased exponentially according to a predefined schedule to ensure progressively selective acceptance of solutions. The genetic algorithm ended after a fixed number of generations (e.g. 100-500).

#### 8.8 Implementation and parameters

The genetic algorithm was implemented in Python, exploiting PyTorch for fitness calculations based on deep learning and standard scientific libraries (e.g. NumPy, Pandas, Matplotlib). Key parameters were optimized through experimental tests, including:

- Population size: typically, 100-500 sequences.
- Generations: 50-500 generations.
- Mutation rates: 0.01 to 0.5 per codon.
- Initial temperature: typically set at 5.0.
- Tournament size: 3 individuals per selection round.

### 9. Analysis of intracellular translatability of GA-modified sequences

#### 9.1 Enzymatic capping and polyadenylation of 100% N1-Methylpseudouridine-5’-Triphosphate(m1y) modified mRNA

To test RNA translatability, batch IVT was performed with 100% substitution of m1y (Trilink N-1081). Subsequently, these purified RNA were capped and polyadenylated using Faustovirus Capping Enzyme (NEB) and *E. coli* Poly(A) Polymerase (NEB) respectively according to manufacturer’s protocols.

#### 9.2 Cell transfection to assess protein expression by ELISA and immunoblotting analysis

Human 293T cells were maintained in Dulbecco’s Modified Eagle Medium supplemented with 10% fetal bovine serum and 1X Penicillin-Streptomycin (Gibco). To test translatability, capped mRNA that has a polyadenylated tail and 100% m1y was transfected using messangerMax (Invitrogen) according to manufacturer’s instructions. For SARS-CoV2 XBB.8, after 72h post-transfection, supernatants were collected, and spike protein concentration determined by ELISA (GeneTex GTX536267) following the manufacturer’s protocol. For hSpCas9, after 72h post-transfection, cells were lysed using lysis buffer (1% NP40, 150mM NaCl, and 25mM Tris-HCl pH7) supplemented with protease inhibitor. Subsequently, the pellet was cleared by centrifugation.

Clear cell lysates were mixed with 1X SDS loading dye (0.8% SDS, 50mM Tris buffer pH6.8, 6% glycerol, 2% b-ME) and resolved by SDS-PAGE. Next, protein samples were transferred to methanol-activated PVDF membranes. Transferred membranes were incubated with relevant antibodies at 4°C overnight: FLAG M2 (Sigma F1804) and GAPDH (Prointech 60004-1-Ig). The next day, after washing three times with Tris-buffered saline with 0.1% Tween 20 (TBST), blots were incubated with horseradish peroxidase-conjugated secondary antibodies. Chemiluminescence was detected using a Bio-Rad ChemiDoc machine.Band intensity was determined by densitometry analysis.

### 10. Analysis of human transcriptome and co-optimization for transcriptional efficiency and pseudoyield

#### 10.1 Prediction of translational efficiency

Translation Efficiency (TE) was predicted using RiboNN^44^. The software was downloaded from https://github.com/Sanofi-Public/RiboNN, and used with the publicly available human model checkpoints and default parameter settings. For each sequence, the TE score was computed as the mean across all available tissue-specific models.

#### 10.2 Analysis of the Human Transcriptome

Human CDS transcript sequences (GRCh38.115) were extracted from the reference FASTA and annotated with the corresponding Ensembl GTF file. For each protein-coding gene, the transcript tagged as “canonical” or “MANE Select” was selected as the primary isoform, and the remaining transcripts were discarded as alternative isoforms.

Once the human transcriptome was obtained, to include 19761 transcripts, we predicted TE using RiboNN as the average across all available tissues, and PY using MAP-Net as the consensus across all manufacturing methods.

Initiation and elongation rates for all transcripts were extracted from previously published results^43^.

#### 10.3 Co-optimization for TE and PY

We adapted our previously described genetic algorithm (GA) (see Section 8) to perform multi-objective optimization, simultaneously maximizing both TE and PY. In this modified GA, the RiboNN-predicted TE serves as a second fitness component alongside the MAP-Net PY score.

To jointly optimize the two objectives, we employed a Pareto optimization scheme, where individuals are ranked based on non-dominance rather than on a single aggregated score. Under this framework, a solution is favored if it improves at least one objective without substantially worsening the other. We assigned equal importance to TE and pseudo-yield when constructing the Pareto front, ensuring that the evolutionary pressure does not bias the search toward a single objective.

#### 10.4 Codon optimization of the Wuhan Spike protein

The cDNA coding sequence for the Spike ORF of the SARS-COV-2 Wuhan-Hu-1 complete genome was downloaded from NCBI RefSeq NC_045512.2 entry (https://www.ncbi.nlm.nih.gov/nuccore/1798174254) at coordinates: 21563..25384. We introduced K986P and V987P mutations to stabilize spike protein in its prefusion state^62^ and defined this sequence as “wild-type” (WT).

The following software programs were used to codon-optimize the WT sequence:

- **GenSmart**: https://www.genscript.com/gensmart-free-gene-codon-optimization.html?src=google&jiraid=12194&gad_campaignid=10424380693
- **Twist**: https://www.twistbioscience.com/resources/digital-tools/codon-optimization-tool?adgroup=166838088849&creative=747091457754&device=c&matchtype=p&location=9062519&gad_campaignid=21430349275
- **IDT**: https://sg.idtdna.com/pages/tools/codon-optimization-tool?gad_campaignid=2010842806
- **VB**: https://en.vectorbuilder.com/tool/codon-optimization.html
- **ExpO**: https://www.novoprolabs.com/tools/codon-optimization
- **GENEWIZ**: https://www.genewiz.com/public/services/gene-synthesis/codon-optimization
- **Optimzer**: http://genomes.urv.es/OPTIMIZER/

When the software requested to indicate a codon-usage target, we provided the *Homo sapiens* codon usage data downloaded from https://www.kazusa.or.jp/codon/cgi-bin/showcodon.cgi?species=9606:

**Homo sapiens [gbpri]: 93487 CDS’s (40662582 codons)**

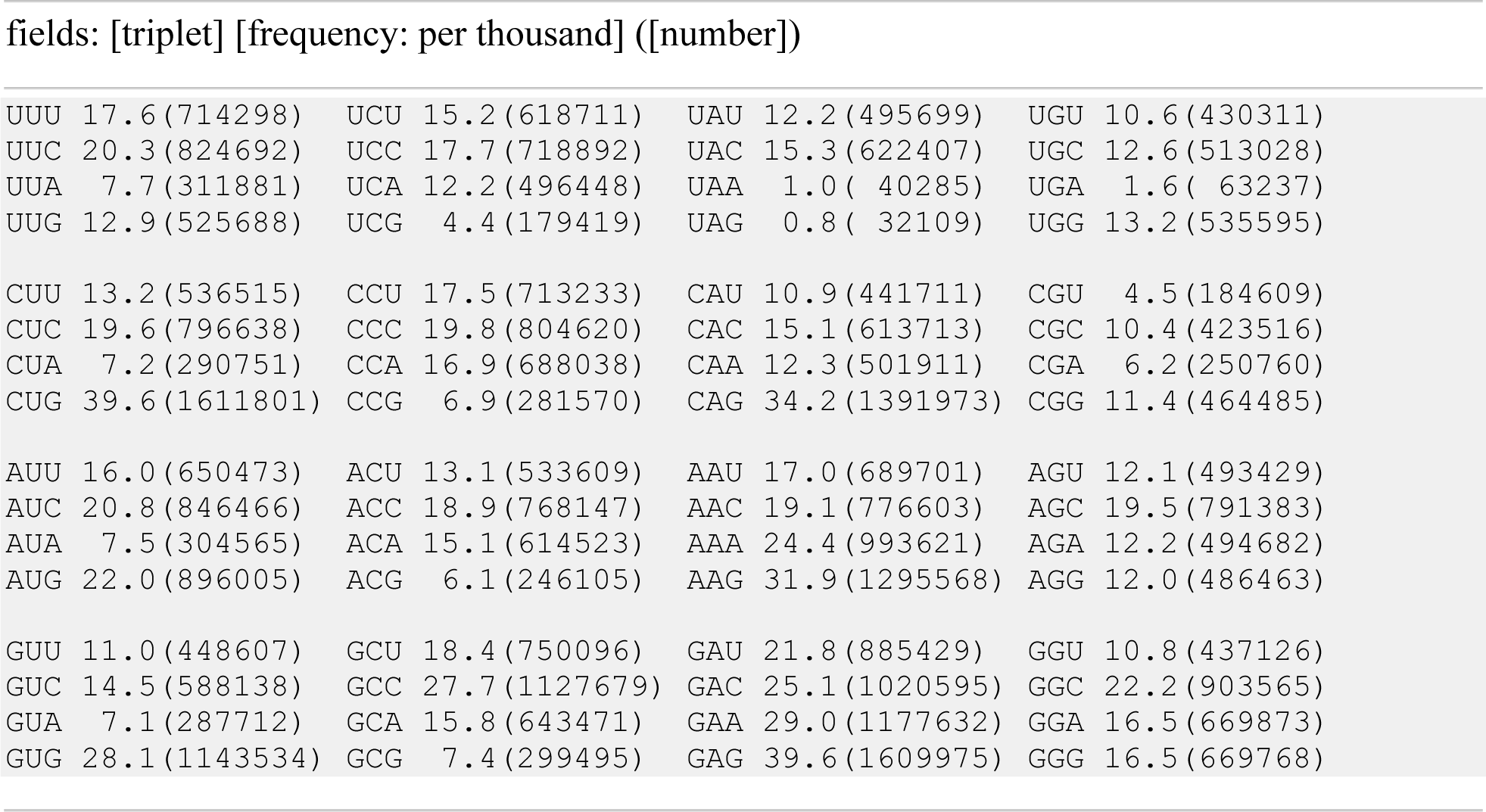

Each software was run three times independently. If the different runs produced the same output, only one sequence was stored and analyzed. When the three runs produced different outputs, all the generated sequences were stored and analyzed.

The sequences for the commercial BioNTech and Moderna vaccine products were downloaded from:

- BNT162b2: https://www.ncbi.nlm.nih.gov/nucleotide/PP544446.1?report=genbank&log$=nuclalign&blast_rank=1&RID=BXW4B3KJ014
- mRNA1273: https://www.ncbi.nlm.nih.gov/nucleotide/OK120841.1?report=genbank&log$=nuclalign&blast_rank=2&RID=BXW59AJA014

As described in 10.3, a MAP-Net- and RiboNN-based GA was used to co-optimize the WT simultaneously sequence for TE and PY. Three different sequences were selected from the population of the last generation of the GA, with highest TE and PY.

All the sequences are reported in Table S1e.

## Extended Data

**Figure S1.**
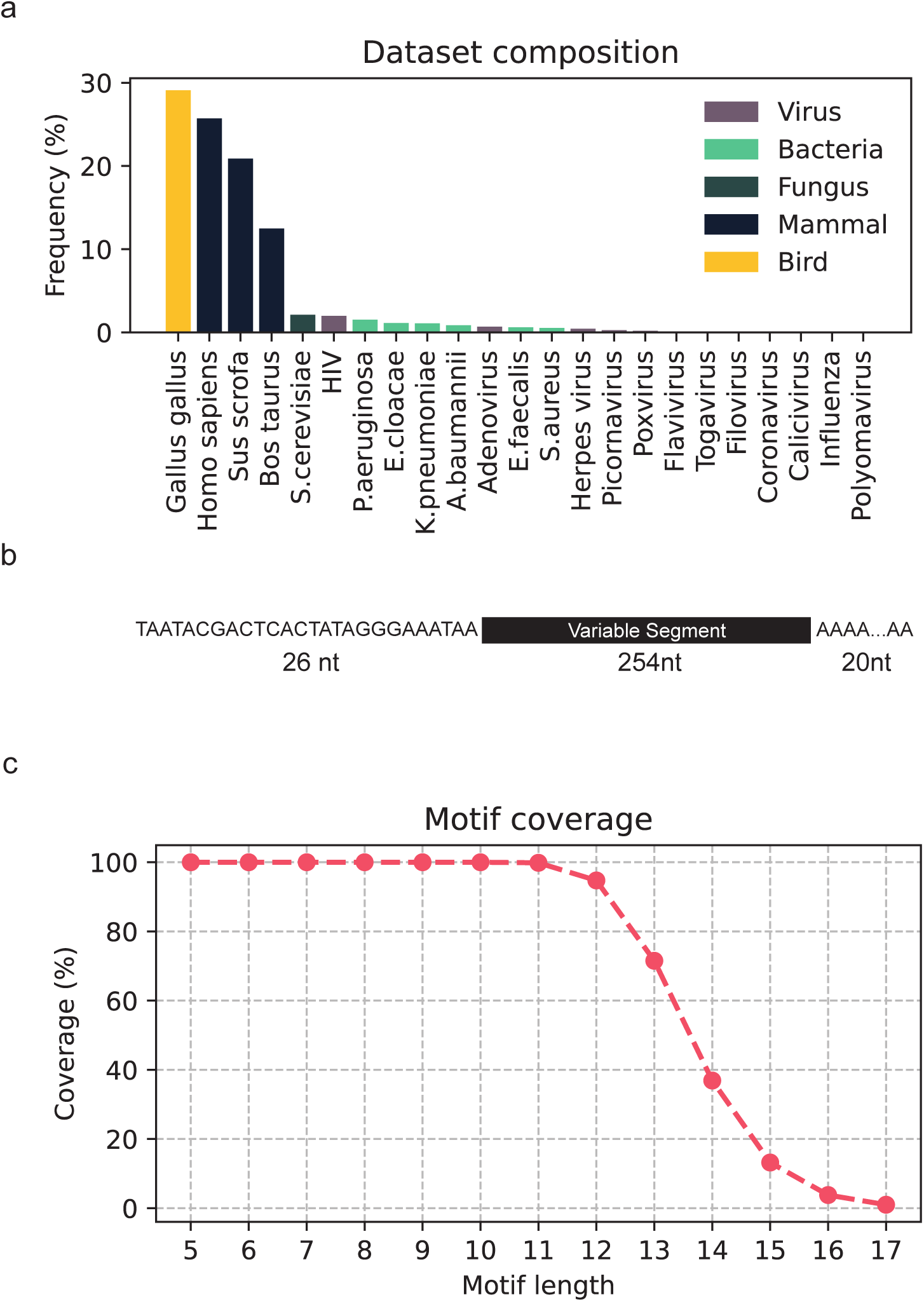
Composition of the 1 Million DNA library. The library was constructed using tiled genomic fragments from different classes of organisms to represent a space of likely candidate RNA of interest for manufacturing, including eukaryotic, bacterial, fungal and viral RNAs**. A.** Percentage of genomic origin of fragments per species. **B.** DNA template structure: each template uses a common 26nt IVT initiation sequence at 5’ and a 20nt poly-A tail at 3’, which frame a 254nt variable fragment extracted from coding transcripts**. C.** Representation of k-mers in the template library, for various motif lengths k.

**Figure S2.**
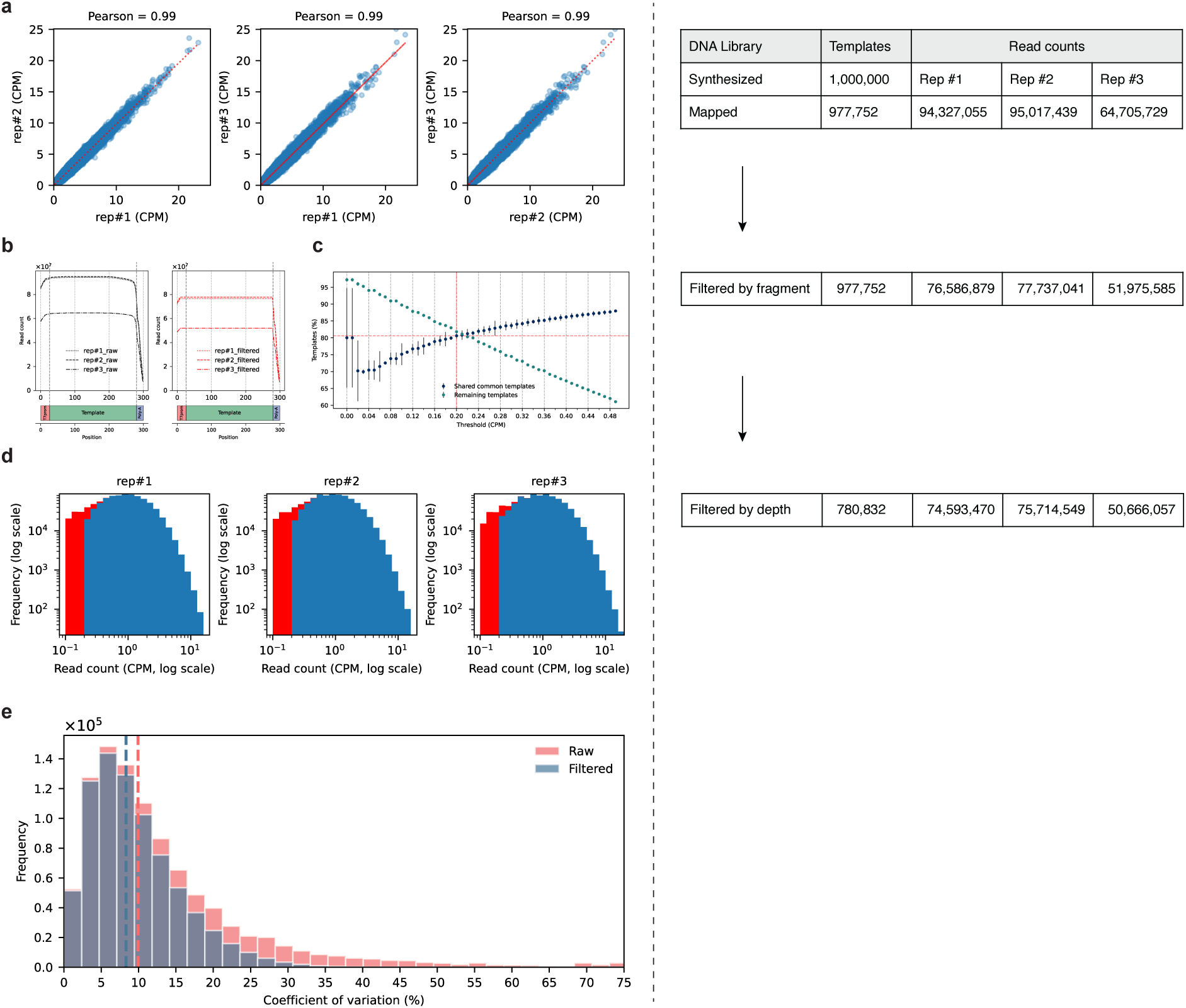
DNA library sequencing and quantification. **A.** A library of 1 million diverse DNA templates was sequenced in triplicate using Oxford Nanopore sequencing. Scatter plots compare counts per million (CPM) between replicates; red lines represent linear regressions**. B.** Total sequencing depth across all 1 M templates, shown before (left) and after (right) coverage filtering. Only reads spanning the library’s variable region were retained. **C.** Effect of CPM-threshold filtering on template retention. For thresholds from 0 to 0.5, blue points indicate the number of templates shared across all replicates with CPM below each threshold; green points show the total templates remaining after filtering. The dashed red line marks the chosen cutoff (CPM = 0.2), which retains 80 % of templates**. D.** Distribution of CPM values in each replicate, before (red) and after (blue) removing templates with CPM < 0.2**. E.** Frequency distribution of the coefficient of variation (CV) for CPM among the 3 replicates, calculated before (red) and after (blue) filtering by depth. Dotted lines indicate the respective median CVs.

**Figure S3.**
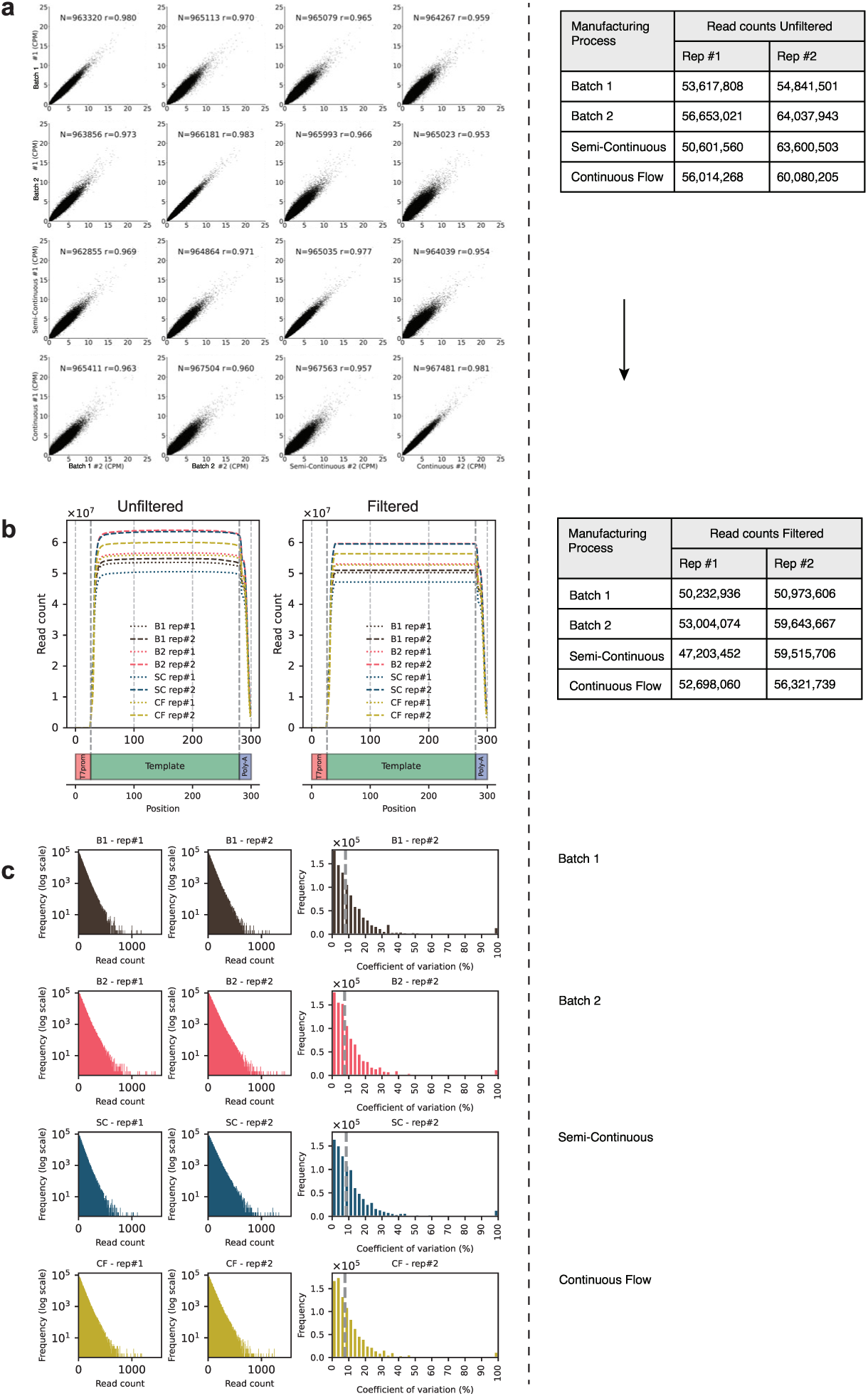
IVT-mRNA library sequencing and quantification. **A.** Pairwise correlation of normalized direct RNA sequencing read counts (counts per million reads, CPM) from IVT-RNA pools generated with the same IVT process (diagonal) and different IVT processes (off-diagonal**). B.** IVT total sequencing depth across all 1 M templates, shown before (left) and after (right) coverage filtering. Only reads spanning the library’s variable region were retained. **C.** For each IVT process (Batch using commercial kit = brown, Batch using in-house materials= red, Semi-continuous = blue, Continuous = yellow) shown are the distribution of CPM for each replicate (left), and the distribution of the coefficient of variation (CV) calculated for each template between replicates (right). The dotted line represents the median CV value.

**Figure S4.**
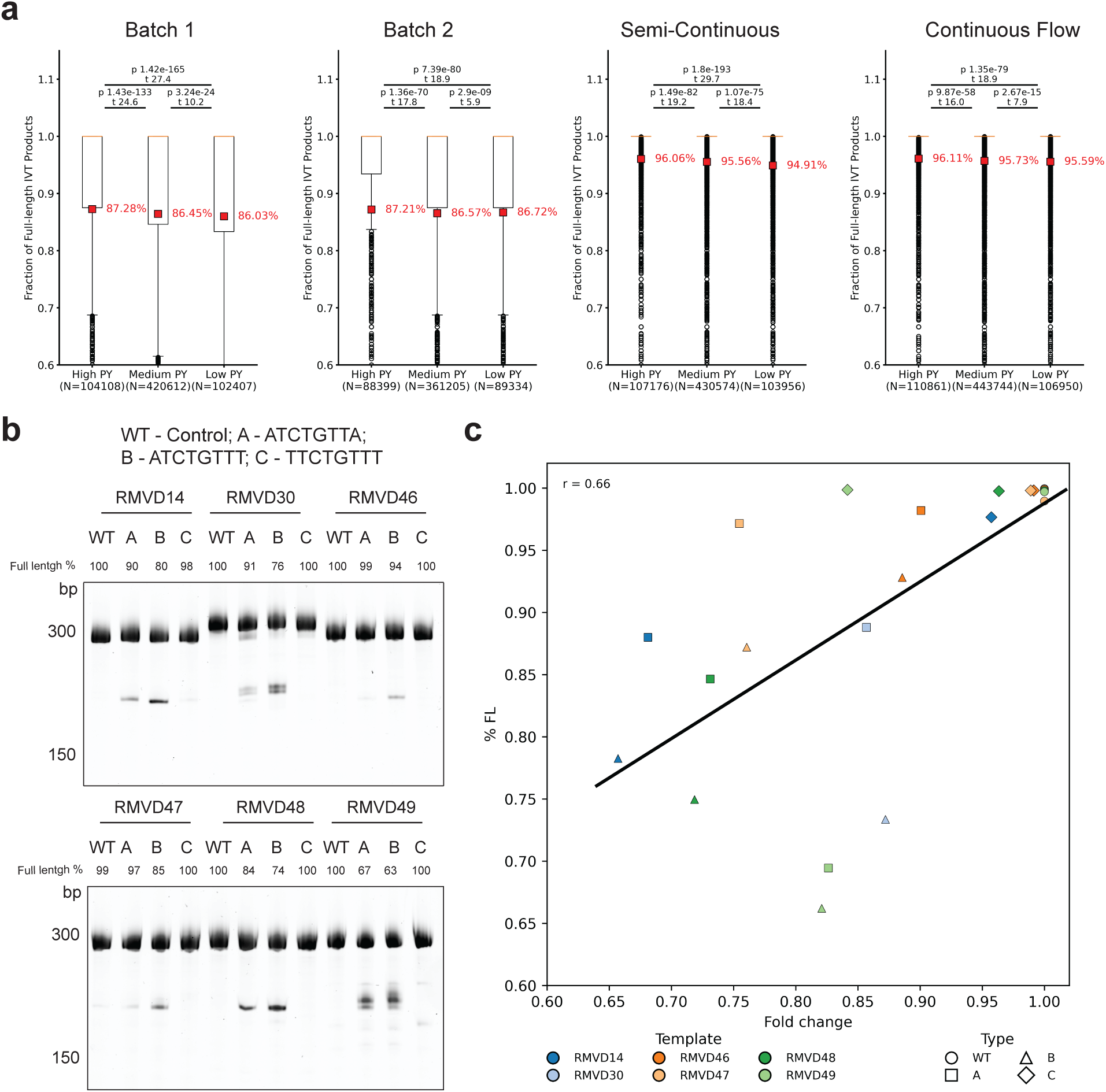
Characterization of IVT termination. **A.** Distributions of full-length transcript percentages in high PY, medium PY, and low PY templates from various manufacturing processes are shown. N represents the number of templates in each PY category and red numbers indicate the mean value in each distribution. p indicates P-value and t is the Mann-Whitney U-test statistic between the 3 distributions**. B.** RMVD DNA templates containing WT (no motif control), A (ATCTGTTA), B (ATCTGTTT), and C motifs (TTCTGTTT) were used for batch IVT. After purification, 50ng IVT products were analysed by UREA-TBE-PAGE and the percentage of full-transcript was calculated by dividing the band intensity of full-length transcript by the sum of band intensity of full-length transcript and truncated products. The band intensity was determined by densitometry analysis. **C.** Correlation between experimental results of full-length percentage (y-axis) versus predicted fold change of PY (x-axis). This fold change in PY (relative to WT within respective RMVD contexts) represents the percentage of full-length as predicted PY accounts for full-length transcript only.

**Figure S5.**
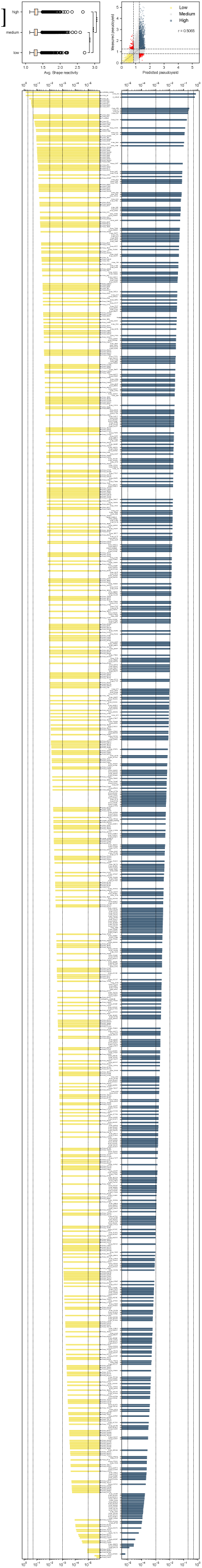
Features differentiating low- *vs.* high-pseudoyield sequences. **A.** Box plot of the SHAPE-MaP secondary structure stratified by PY class. Medians are shown in orange, black boxes represent interquartile ranges (IQRs), whiskers are 1.5*IQR, outliers as circles. Horizontal brackets denote significant differences between low- *vs.* high- and medium-*vs.* low- (p < 0.01**). B.** scatterplot of predicted *vs.* measured PYs for templates in the test sets. Templates with concordant predicted and measured high/low PYs are shown in blue/yellow, respectively. Templates with predicted and/or measured medium PY are gray. Templates with discordant predicted and measured low/high PYs are shown in red. The solid line is a linear regression of predicted *vs.* measured PYs. **C.** LASSO feature coefficients for the 906 predictors. Yellow bars correspond to features that drive the low-PY class; blue bars correspond to features that drive the high-PY class.

**Figure S6.**
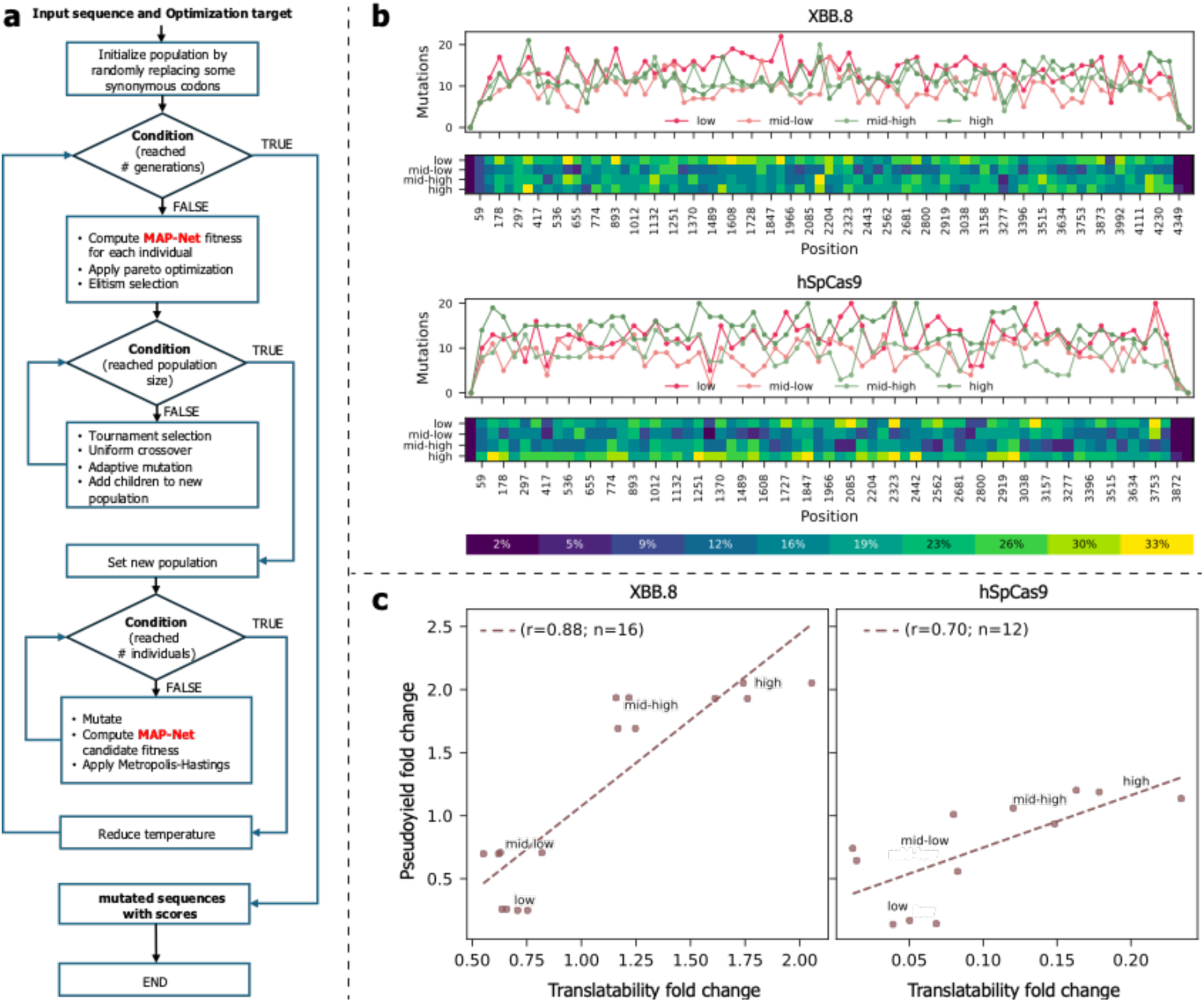
Application of MAP-Net to natural genes. **A.** Structure of the genetic algorithm (GA) reading a DNA sequence in input, using MAP-Net (in red) as a fitness criterion to produce synonymously mutated DNA sequences, along with predicted PY as the associated score. **B.** Distribution of mutations applied to XBB.8 (top panel) and hSpCas9 (bottom panel). The heatmaps represent the percentage of mutated residues for each 60nt-long bin, while the line graph shows the absolute number of mutations introduced. c) Scatter plots comparing the full-length IVT product yields to the sequence translatability. Both values are reported as a fold change *vs.* the WT sequence. The solid line is a linear regression of translatability *vs.* measured yield.

