## Supplementary material for "A Universal, AI-based Design Framework for Efficient Manufacturing of mRNA Therapeutics": All Supplemental Table Legends, Supplemental Tables S2, S3, S5

**The PDF file includes:**

Tables S1 to S7

**Table S1**: see file “Supplementary Table 1 - sequences.xlsx”

a) Primers used in this study.
b) Individual Validation: Template sequences used for experiments in Figure 2D.
c) Termination Validation: Template sequences used for experiments in Figure S4B.
d) Gene Validation: Template sequences used for experiments in Figure 5.
e) Spike Wuhan-Hu-1 co-optimization: sequences used in Figure 6e

| **ID** | **Sample** | **IVT Protocol** | **Technology** |
| --- | --- | --- | --- |
| PAS52745 | DNA | - | ONT |
| PAU02392 | DNA | - | ONT |
| PAO26582 | DNA | - | ONT |
| RHX892 | DNA | - | Illumina |
| PAQ17590 | IVT-mRNA | Batch (commercial kit) | ONT |
| PAS86168 | IVT-mRNA | Batch (commercial kit) | ONT |
| PAS86151 | IVT-mRNA | Continuous | ONT |
| PAU02059 | IVT-mRNA | Continuous | ONT |
| PAS86136 | IVT-mRNA | Semi-continuous | ONT |
| PAS86096 | IVT-mRNA | Semi-continuous | ONT |
| PAS92305 | IVT-mRNA | Batch (in-house) | ONT |
| PAS92258 | IVT-mRNA | Batch (in-house) | ONT |
| RHX1129 | IVT-mRNA | Batch (commercial kit) | Illumina |
| RHX1130 | IVT-mRNA | Batch (commercial kit) | Illumina |
| RHX1131 | IVT-mRNA | Batch (commercial kit) | Illumina |
| RHX1151 | IVT-mRNA | Batch (in-house) | Illumina |
| RHX1152 | IVT-mRNA | Batch (in-house) | Illumina |
| RHX1153 | IVT-mRNA | Batch (in-house) | Illumina |
| RHX1154 | IVT-mRNA | Continuous | Illumina |
| RHX1155 | IVT-mRNA | Continuous | Illumina |
| RHX1156 | IVT-mRNA | Continuous | Illumina |
| RHX1287 | IVT-mRNA | Semi-continuous | Illumina |
| RHX1288 | IVT-mRNA | Semi-continuous | Illumina |
| RHX1289 | IVT-mRNA | Semi-continuous | Illumina |

**Table S2**. Accession numbers and description of next-generation sequencing runs. Datasets for are available for download at [links to be provided]

| **Name** | **# of features** | **Description** |
| --- | --- | --- |
| nucleotide_composition | 4 | Fraction (or count) of each nucleotide (A, C, G, T) in the sequence |
| pseudo_nucleotide_composition | 4+1(θ) | Computes standard nucleotide composition, then adjusts each by a sequence‐order term θ^56^ |
| molecular_weight | 1 | Molecular mass of RNA transcript |
| melting_temperature | 1 | Predicted Wallace melting temperature^57^ |
| purine_content | 1 | Fraction of purine bases (A + G) |
| pyrimidine_content | 1 | Fraction of pyrimidine bases (C + T/U) |
| Shannon_entropy | 1 | Shannon entropy^58^ |
| longest_repeated_substring | 1 | Length of the longest exact substring that appears at least twice |
| Kolmogorov_complexity | 1 | Approximate complexity estimated via sequence compression |
| repeat_content_n for n ∈ {2,3,4,5} | 4 | Enumerates all unique n-mers, sums how often each occurs beyond its first appearance, and divides by sequence length |
| n_gram_entropy for n ∈ {2,3,4,5} | 4 | Counts all overlapping n-mers, converts counts to probabilities, and computes Shannon entropy |
| 2_mer_composition | 16 | Fraction of each dinucleotide |
| 3_mer_composition | 64 | Fraction of each trinucleotide |
| 4_mer_composition | 256 | Fraction of each tetranucleotide |
| 5_mer_composition | 1024 | Fraction of each pentanucleotide |

**Table S3**. DNA-encoded features for LASSO analysis

**Table S4**: See file “Supplementary Table 4 - feat_selection.xlsx”

Legend: Statistical analysis of primary sequence features. For each feature, a non-parametric Mann-Whitney U test was conducted to compare two template groups: 'high-pseudoyield' and 'low-pseudoyield'. This analysis generated a U statistic, a p-value, and an adjusted p-value. Additionally, odds ratios were calculated. The analysis also includes the number of templates analyzed in each group, along with basic statistics such as the median and average for each group.

| **feature_1** | **feature_2** | **Pearson’s correlation** |
| --- | --- | --- |
| melting_temperature | ~~cg_content~~ | 1 |
| melting_temperature | ~~at_content~~ | 1 |
| ~~at_content~~ | ~~cg_content~~ | 1 |
| pyrimidine_content | ~~purine_content~~ | 1 |
| ~~Pseudo_C_content~~ | ~~C_content~~ | 1 |
| ~~Pseudo_T_content~~ | ~~T_content~~ | 1 |
| G_content | ~~Pseudo_G_content~~ | 1 |
| A_content | ~~Pseudo_A_content~~ | 1 |
| ~~7-mer_entropy~~ | ~~9-mer_entropy~~ | 0.974 |
| 5~~-mer_TAGGG~~ | 5-mer_GTTAG | 0.967 |
| 5-mer_entropy | ~~repeat_content_5~~ | 0.962 |
| ~~5-mer_GTTAG~~ | ~~5-mer_AGGGT~~ | 0.961 |
| 5-mer_TTAGG | 5-mer_~~GTTAG~~ | 0.958 |
| 5-mer_TTAGG | ~~5-mer_AGGGT~~ | 0.946 |
| ~~5-mer_TAGGG~~ | ~~5-mer_AGGGT~~ | 0.939 |
| ~~3-mer_TTT~~ | ~~2-mer_TT~~ | 0.935 |
| 5-mer_TTAGG | ~~5-mer_TAGGG~~ | 0.934 |
| ~~2-mer_TT~~ | ~~Pseudo_T_content~~ | 0.93 |
| ~~2-mer_TT~~ | ~~T_content~~ | 0.93 |
| ~~5-mer_GGTTA~~ | ~~5-mer_AGGGT~~ | 0.927 |
| 5-mer_GGGTT | ~~5-mer_GGTTA~~ | 0.925 |
| ~~5-mer_TAGGG~~ | ~~5-mer_GGTTA~~ | 0.925 |
| ~~5-mer_GGTTA~~ | ~~5-mer_GTTAG~~ | 0.924 |
| 3-mer_entropy | ~~repeat_content_4~~ | 0.922 |
| ~~repeat_content_5~~ | repeat_content_4 | 0.918 |
| 5-mer_TTAGG | ~~5-mer_GGTTA~~ | 0.916 |
| 4-mer_TTTT | ~~3-mer_TTT~~ | 0.915 |
| 2-mer_CC | ~~Pseudo_C_content~~ | 0.914 |
| 2-mer_CC | ~~C_content~~ | 0.914 |
| 5-mer_CACAC | ~~5-mer_ACACA~~ | 0.911 |
| 5-mer_CCCTA | ~~5-mer_CTAAC~~ | 0.911 |
| 5-mer_entropy | ~~7-mer_entropy~~ | 0.907 |
| ~~2-mer_AA~~ | ~~3-mer_AAA~~ | 0.905 |
| ~~Pseudo_A_content~~ | 2-mer_AA | 0.904 |
| A_content | ~~2-mer_AA~~ | 0.904 |
| 4-mer_TTTT | ~~5-mer_TTTTT~~ | 0.901 |

**Table S5.** Redundant features originated from LASSO correlation analysis. Removed features are strike through

**Table S6**: see file “Supplementary Table 6 - ssearch.xlsx”. The 1M template library was searched for the presence of known T7 terminator sequences using SSEARCH. The table includes all match results.

**Table S7:** see file “Supplementary table 7 - motif termination strength.xlsx”

Fit parameters of Equation 11 to all 6-mer motifs for all Illumina samples. Individual fit parameters λ, μ, and s as well as the mixture parameters A and B are given. Termination strength is reported as B/A, with higher values indicating sigmoid termination behavior as opposed to stochastic decay.
